# Clustering and reverse transcription of HIV-1 genomes in nuclear niches of macrophages

**DOI:** 10.1101/2020.04.12.038067

**Authors:** Elena Rensen, Florian Mueller, Viviana Scoca, Jyotsana J. Parmar, Philippe Souque, Christophe Zimmer, Francesca Di Nunzio

## Abstract

In order to replicate, the Human Immunodeficiency Virus (HIV-1) reverse transcribes its RNA genome into DNA, which subsequently integrates into host cell chromosomes. These two key events of the viral life cycle are commonly viewed as separate not only in time but also in cellular space, since reverse transcription (RT) is thought to be completed in the cytoplasm before nuclear import and integration. However, the spatiotemporal organization of the early replication cycle in macrophages, natural non-dividing target cells that constitute reservoirs of HIV-1 and an obstacle to curing AIDS, remains unclear. Here, we demonstrate that infected macrophages display large nuclear foci of viral DNA and viral RNA, in which multiple genomes cluster together. These clusters form in the absence of chromosomal integration, sequester the paraspeckle protein CPSF6 and localize to nuclear speckles. Strikingly, we show that viral RNA clusters consist mostly of genomic, incoming RNA, both in cells where RT is pharmacologically suppressed and in untreated cells. We demonstrate that, after temporary inhibition, RT can resume in the nucleus and lead to vDNA accumulation in these clusters. We further show that nuclear RT can result in transcription competent viral DNA. These findings change our understanding of the early HIV-1 replication cycle, and may have implications for understanding HIV-1 persistence.

---

Productive infection by the Human Immunodeficiency Virus 1 (HIV-1), the causative agent of AIDS, requires reverse transcription (RT) of the viral RNA (vRNA) genome into double-stranded viral DNA (vDNA) and subsequent integration of the vDNA into host cell chromosomes. Studies in immortalized cell lines such as HeLa and activated CD4+ T cells have established a spatiotemporal sequence of events where: (i) vDNA is synthesized by RT in the cytoplasm, with concomitant degradation of the template vRNA, (ii) the vDNA genome translocates into the nucleus, (iii) the integrase enzyme (IN) inserts the vDNA into the genome, (iv) the integrated vDNA genome undergoes transcription that leads to viral progeny (Freed, 2001; Hu & Hughes, 2012). How the HIV-1 replication cycle proceeds in other cell types remains comparatively underexplored. Indeed, different cell types can exhibit widely divergent responses to viral attacks, owing to e.g. different restriction factors or immune cell defense mechanisms (Stremlau et al, 2004; Goujon et al, 2013; Lahaye et al, 2018; Rasaiyaah et al, 2013). Macrophages are terminally differentiated, non-dividing cells derived from blood monocytes, which play a critical role in the innate and adaptive immune response (Koppensteiner et al, 2012; Gordon & Taylor, 2005; Ganor et al, 2019). Along with activated CD4+ T cells, macrophages are natural target cells for HIV-1 and accumulating evidence points to a critical role of these cells in viral persistence, which remains a major roadblock to eradicating HIV (Ganor et al, 2019; Honeycutt et al, 2017). Despite this importance, the early steps of HIV-1 infection in macrophages remain elusive. Here, we use imaging approaches to visualize and quantify the cellular localizations of vDNA and vRNA in infected macrophages and study their link with RT. Our data reveal that genomic vRNA forms nuclear clusters that associate with nuclear speckle factors, and provide surprising evidence that these structures can harbor a nuclear RT activity.

## HIV-1 genomes form large nuclear foci in infected macrophages

To study HIV infection in macrophages, we used primarily ThP1 cells, a human monocytic cell line, and differentiated them into macrophage-like cells by stimulation with phorbol esters (Schwende et al, 1996). We infected these cells with VSV-G pseudotyped HIV-1 carrying the HIV-2 accessory protein Vpx (unless stated otherwise), which overcomes the natural resistance of macrophages to viral infection by counteracting the host restriction factor SAMHD1 (Laguette et al, 2011). To enable fluorescent labeling of the virus, we used a virus carrying the endogenous viral integrase (IN) gene fused to an HA-tag (Petit et al, 2000, 1999). The tagged virus is similarly infectious as the untagged virus (Petit et al, 1999; Blanco-Rodriguez et al, 2020). We analyzed reverse transcription (RT) with qPCR by measuring viral DNA synthesis and the presence of nuclear viral DNA (vDNA) forms including 2LTRs at different times post infection (p.i.). In parallel, we also measured the number of integrated proviruses, revealing a low level of integration in these cells at 24 h p.i. (**Fig. S1**). These assays indicated that RT peaks at ~24 h p.i., and that the formation of episomal nuclear forms (2LTRs), peaks at ~30 h p.i., confirming that these early steps of HIV-1 infection in ThP1 macrophage-like cells are delayed relative to HeLa cells (Arfi et al, 2008).

To visualize the reverse transcribed viral DNA genome, we infected ThP1 cells in the presence of the nucleotide analog EdU for 24 hours (Peng et al, 2014; Stultz et al, 2017) with a multiplicity of infection (MOI) of 50 (unless otherwise stated), as measured by qPCR on 293T cells. Fluorescent visualization of EdU was performed at 48 h p.i. by click chemistry, unless stated otherwise. Because the vast majority of ThP1 cells were terminally differentiated, EdU did not incorporate into host cell chromosomes. A minority of cells (~5 %) exhibited bright EdU signal throughout the nucleus, clearly indicating that these cells failed to differentiate and continued to replicate their DNA (**Fig. S2A**). We excluded these cells from further analysis throughout this study and only considered the remaining, terminally differentiated cells. Our images revealed strikingly large and bright EdU foci in the nuclei of infected cells, whereas uninfected control cells displayed only a very weak background signal (**Fig. 1A, Movie S1, Fig. S2B-C**). While the nuclear envelope of some cells displayed invaginations, dual color imaging of EdU with immunostained lamin in infected cells confirmed that these foci are located within the nuclear lumen (**Fig. 1C, Fig. EV1**). Quantifications indicated that ~73% of cells contained one or more foci (**Fig. 1J**) and that foci had a median size (as measured by the full width at half maximum, FWHM, of intensity profiles) of ~600 nm (interquartile range ~175 nm; n=40 foci) (**Fig. 1B,F**). In addition to these large foci, some infected cells also showed discrete nuclear punctae of much lower brightness, with a size (FWHM) of ~370 nm, close to the microscope’s theoretical resolution of ~300 nm and hence consistent with particles of similar or smaller size (**Fig. 1A,F**). Triple color imaging of EdU together with immunolabeled capsid (CA) and integrase (IN) exhibited clear colocalization, thereby simultaneously confirming the viral nature of EdU-labeled DNA, and showing that the EdU foci (hereafter referred to as ‘vDNA foci’) are enriched in these viral proteins (**Fig. 1D, Fig. S3**). Importantly, although our experiments used viral particles incorporating Vpx to increase HIV-1 infection efficiency, we observed that in absence of Vpx, infected ThP1 cells also form nuclear vDNA foci, ruling out the possibility that these foci are an artifact due to the presence of Vpx (**Fig. S4**).

**Figure 1:**
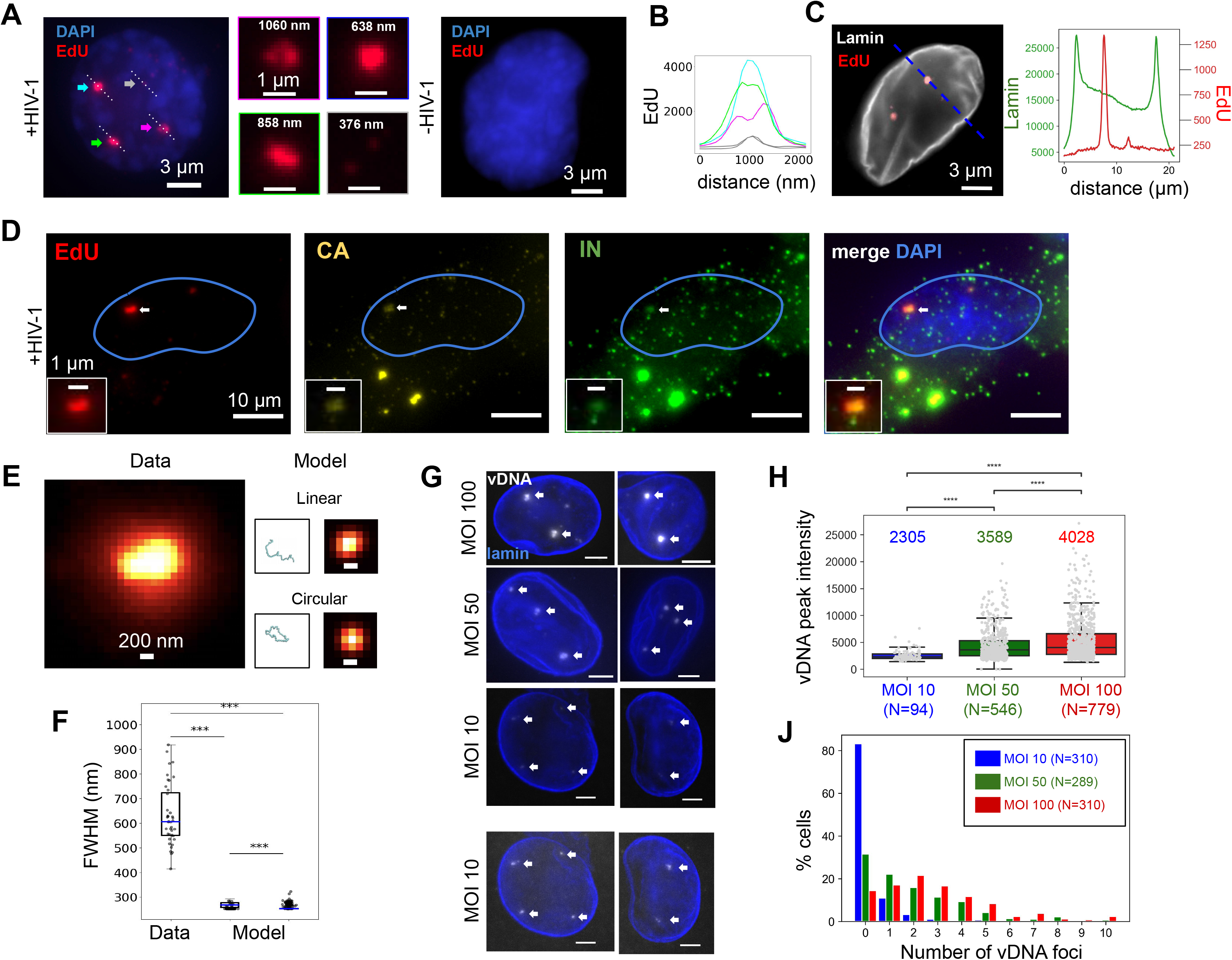
HIV genomes form large nuclear foci in macrophages. (**A-C**) EdU-labeled viral DNA forms large foci in ThP1 cells infected by HIV-1. EdU images are shown in red, DAPI images in blue. (**A**) Left: nucleus of an infected cell at 48 h p.i. displays three large and bright EdU foci (colored arrows), as well as small and dim punctae (grey arrows). Center: Magnified views have colored borders matching the colors of the arrows. 3D images show that EdU foci are located within the nucleoplasm (**Movie S1, Fig. S2**). Right: nucleus of an uninfected cell displays no EdU signal. (**B**) Graph shows EdU intensity profiles along lines shown in **A**, with colors matching the corresponding arrows. (**C**) Dual-color image of an infected cell with EdU (red) and immunolabeled lamin (grey). Curves to the right plot the intensity profile of EdU and lamin along the dashed blue line. The EdU intensity peaks do not coincide with lamin enrichment. See also **Fig. EV1**. (**D**) Multi-color images of an infected cell showing EdU (red) with CA (yellow) and integrase (green). The colocalization of nuclear EdU foci with CA and IN confirms that EdU specifically labels viral DNA and shows the presence of these proteins in vDNA foci. See also **Fig. S3**. (**E,F**) vDNA foci are much larger than the predicted size of single viral genomes. (**E**) Left image shows an observed, EdU-labeled vDNA focus. Right: simulations of a single linear or circular 10 Kb long chromatinized DNA polymer chain and corresponding predicted images in diffraction-limited (~300 nm resolution) microscopy. (**F**) Boxplots show the distribution of sizes (FWHM) of n=40 measured vDNA foci (for MOI 100) compared to the sizes predicted for linear (left) and circular (right) chains based on n=100 simulated configurations each. All differences are highly significant (Wilcoxon test data vs. model: *p* ≈ 3 · 10^-20^, circular vs. linear model: *p* ≈ 10^-17^). (**G**) Images of vDNA foci in ThP1 cells infected with multiplicities of infection (MOI) 10, 50 and 100. Images for MOI 10 are shown at two different contrast levels to reveal dimmer spots. (**H**) Boxplots compare the peak intensities inside individual vDNA foci for MOIs 10, 50 and 100. Intensities increase significantly with MOI (Wilcoxon tests: MOI 10 vs. MOI 50: p ≈ 5 · 10^-16^; MOI 50 vs. MOI 100: *p* ≈ 5 · 10^-5^; MOI 10 vs. MOI 100: *p* ≈ 2 · 10^-23^). (**J**) Histograms show the number of foci per nucleus for MOIs of 10 (blue), 50 (green) and 100 (red).

We next asked if the observed vDNA foci might correspond to individual viral genomes by further analyzing their size and dependence on the MOI. First, we considered the possible size of a single ~10 Kb long piece of DNA, the approximate length of the HIV-1 genome. We used polymer simulations that model the chromatinized vDNA as a linear or circular chain of nucleosomes (Arbona et al, 2017; Geis & Goff, 2019) (**Fig. 1E, Fig. EV2**). Our simulations predict a distribution of apparent sizes for a single 10 Kb long HIV-1 genome that is close to the microscope’s spatial resolution of ~300 nm, hence is much smaller than the observed sizes of nuclear vDNA foci (**Fig. 1F**). Thus, nuclear vDNA foci are larger than expected for single genomes, suggesting that they may comprise multiple genomes. Second, we analyzed the intensity of vDNA foci for MOIs of ~10, ~50 and ~100 (**Fig. 1G,H**). Note that EdU labeling does not involve signal amplification, enabling a quantitative interpretation of the measured intensities. If individual foci correspond to individual genomes, their number is expected to increase proportionally with the MOI, but their intensity should not depend on MOI. In our data, the peak intensity of vDNA foci significantly increased from MOI 10 to MOI 50, and again from MOI 50 to MOI 100 (**Fig. 1H**). Therefore, the MOI dependence of the vDNA intensity also argues for the coexistence of multiple genomes within these foci.

## Nuclear vDNA/vRNA foci contain multiple HIV-1 genomes

To analyze the potential association of vDNA foci with viral RNA (vRNA), we used RNA-FISH probes directed against the POL gene of HIV-1. We verified the specificity of RNA-FISH using uninfected cells, and ruled out the possibility that RNA-FISH probes bind vDNA using infected cells treated with RNase (**Fig. S5A-B**). The dual color EdU/RNA-FISH images showed bright vRNA foci that displayed high and significant colocalization with the vDNA foci (64% of EdU foci contained vRNA, and 30% of vRNA spots contained vDNA; p<0.01), indicating that viral RNA and DNA occupy the same nuclear space (**Fig. 2A,B; Fig. S6**). The vDNA and vRNA intensities in colocalizing spots exhibited a strong positive correlation (**Fig. 2C**).

**Figure 2:**
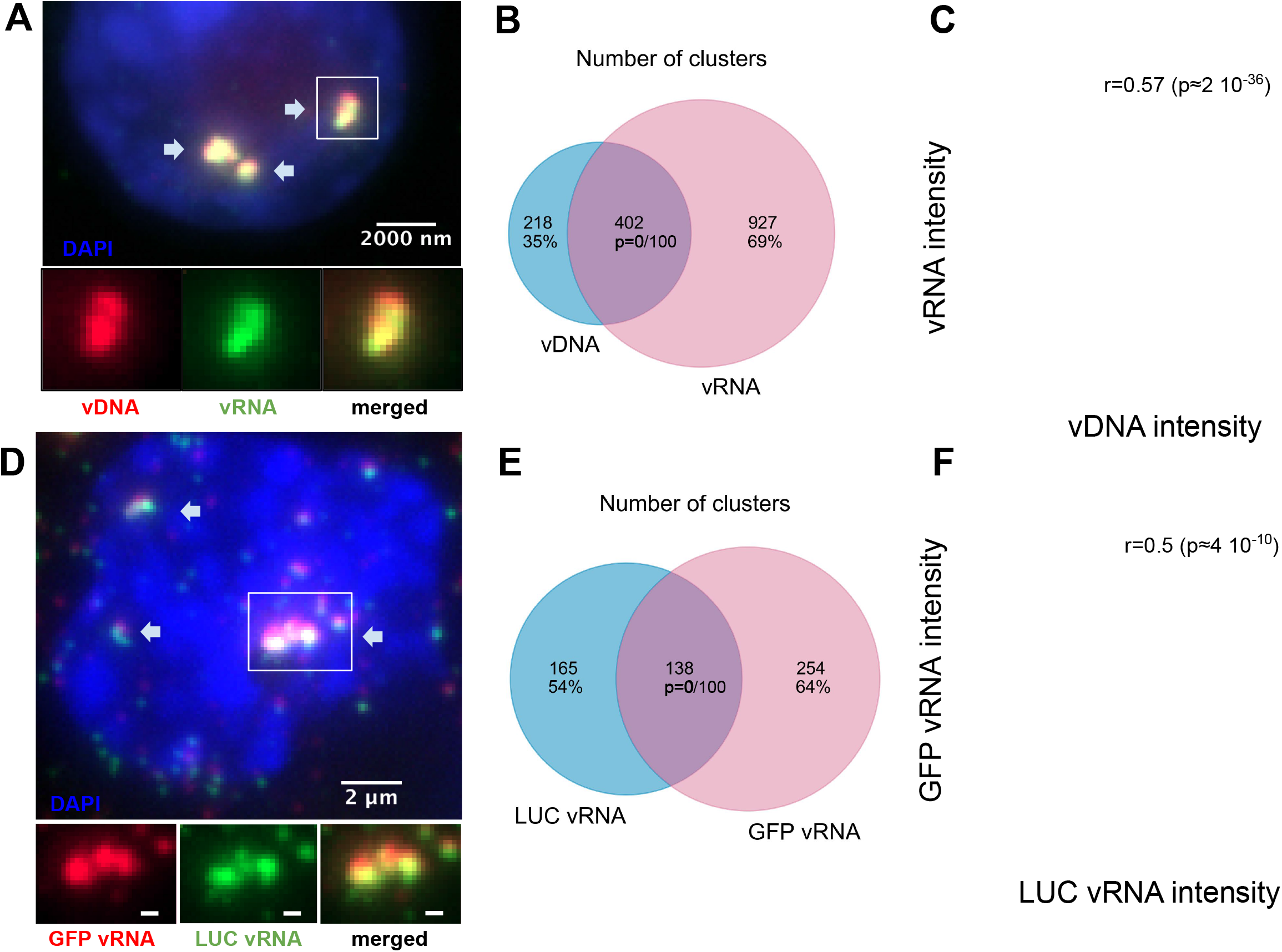
Nuclear foci contain multiple viral RNAs and DNA genomes. (**A-C**) vDNA foci are also foci of vRNA. (**A**) Dual-color image of an infected ThP1 cell showing the viral DNA (red) and the RNA visualized by RNA-FISH (green). See also **Fig. S6**. (**B**) Venn diagram shows the number of vDNA foci and vRNA foci and the number of vDNA foci colocalizing with vRNA foci. The p-value indicates the significance of colocalization based on a jittering analysis (see Methods). (**C**) Scatter plot shows the intensities of vDNA and vRNA in colocalizing foci with the Spearman correlation *r* and associated p-value. (**D,E,F**) In ThP1 cells co-infected with a HIV1-GFP virus and a HIV1-LUC virus, nuclear foci contain mixtures of both viruses. (**D**) Dual-color image shows RNA-FISH against GFP (red) and RNA-FISH against LUC (green). See **Fig. S7**. (**E**) Venn diagram shows the number of GFP RNA foci, the number of LUC RNA foci, and the number of GFP RNA foci colocalizing with LUC RNA foci. (**F**) Scatter plot shows intensities of GFP RNA and LUC RNA in colocalizing foci with the Spearman correlation *r* and associated p-value.

To directly test for the presence of multiple viral genomes in nuclear foci, we coinfected cells with two HIV-1 strains containing different reporter genes, GFP and luciferase (LUC). We then performed dual-color RNA-FISH with two different sets of probes directed against GFP and LUC (**Fig. 2D**). We confirmed the specificity of our RNA-FISH probes using control cells infected with only one of the two reporter viruses (**Fig. S7**). In co-infected cells, RNA-FISH revealed strong colocalization of LUC and GFP RNA (45% of LUC RNA foci contained GFP RNA, and 35% of GFP RNA foci contained LUC RNA; p<0.01) (**Fig. 2E**). In colocalizing foci, intensities of GFP and LUC RNA correlated positively (**Fig. 2F**). Taken together with our observation that vRNA foci occupy the same nuclear regions as vDNA foci (**Fig. 2A,B**), these data firmly establish that nuclear foci are clusters consisting of multiple HIV-1 genomes.

## Viral genome clusters are associated with nuclear body factors

Inspection of our dual EdU/DAPI images indicates that vDNA clusters are located in nuclear regions with lower densities of host cell DNA (**Fig. S8**). This opens the possibility of an association of HIV genomes with nuclear bodies (Spector, 2001; Matera, 1999), membrane-less compartments located in the interchromosomal space. We therefore examined the possible association of viral clusters with host proteins, starting with the cleavage and polyadenylation-specific factor subunit 6 (CPSF6). CPSF6 is known to interact with CA and has been implicated in the regulation of different steps of the viral life cycle from HIV-1 nuclear import to integration site distribution (Price et al, 2012; Achuthan et al, 2018, 6; Buffone et al, 2018; Lee et al, 2010; Burdick et al, 2020). In uninfected cells, CPSF6 displayed a diffuse nucleoplasmic signal; in infected cells, however, CPSF6 accumulated in a small number of nuclear foci, which colocalized with vDNA, in agreement with a recent report (Bejarano et al, 2019) (median Pearson’s r=0.30; Costes p<0.01 for 16 out of 19 regions of interest) (**Fig. 3A,D; Fig. S9**). CPSF6 is known to associate with paraspeckles, nuclear bodies often found in the vicinity of nucleoli and speckles (Fox et al, 2002; Naganuma & Hirose, 2013). To test whether viral clusters associate with paraspeckles, we imaged the non-coding RNA NEAT1 (Nuclear Enriched Abundant Transcript 1/Human nuclear paraspeckle assembly transcript 1), an essential architectural component of paraspeckles (Naganuma & Hirose, 2013; Zhang et al, 2013, 1; Yamazaki et al, 2018, 1). In uninfected cells, NEAT1 was enriched in a small number of nuclear clusters, as expected for paraspeckles (**Fig. 3B; Fig. S10A**). In infected cells, NEAT1 displayed a similar localization pattern, and NEAT1 foci appeared in close vicinity to the vDNA clusters, but did not overlap with them and appeared to be excluded (median Pearson’s r=-0.19; Costes p<0.01 for 19/19 regions of interest) (**Fig. 3B,D; Fig. S10B**). The core paraspeckle protein NONO (p54nrb), recently found to be essential for triggering an immune response to HIV in dendritic cells through interaction with CA (Lahaye et al, 2018), was also excluded from vDNA (**Fig. EV3**). Thus, viral clusters do not localize to paraspeckles, but upon infection, the paraspeckle factor CPSF6 relocates to viral clusters.

**Figure 3:**
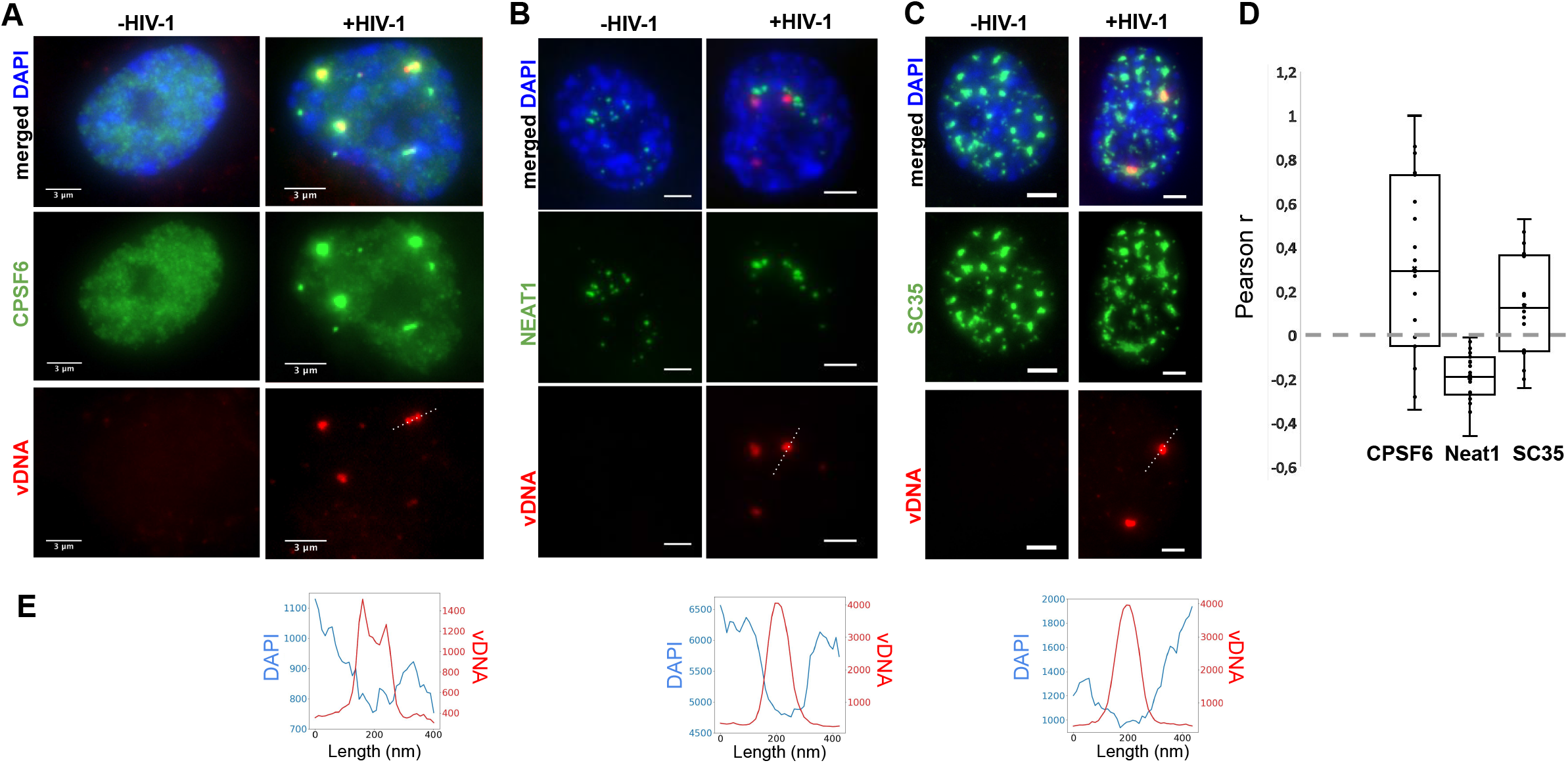
Viral clusters contain specific nuclear body factors. (**A-C**) Images of infected (right) or uninfected (left) ThP1 cells showing the vDNA (EdU) in red, the nucleus (DAPI) in blue, and selected nuclear body factors in green. (**A**) Green image shows immunolabeling of CPSF6. (**B**) Green image shows RNA-FISH against NEAT1. (**C**) Green image shows immunolabeling of SC35. See also **Figs. S9-14**. (**D**) Boxplots show Pearson correlations *r* between vDNA (EdU) and CPSF6, NEAT1 or SC35 in 17-19 regions of interest (ROIs). Significance of positive or negative correlations was assessed using the Costes method of random ROI shifts. Highly significant (p<0.01) positive correlations between vDNA and CPSF6 intensities are found in 16 out of 19 ROIs; highly significant negative correlations between vDNA and NEAT1 are found in 19/19 ROIs, and highly significant positive correlations between vDNA and SC35 are found for 16/17 ROIs. (**E**) Intensity profiles of EdU and DAPI along the dotted lines in the vDNA images above

Paraspeckles are often found in proximity to speckles, nuclear bodies enriched in pre-mRNA splicing factors (Spector & Lamond, 2011). Therefore, our data raised the possibility that viral clusters are associated with speckles. To investigate this, we imaged the non-small nuclear ribonucleoprotein particle factor SC35, a well-studied splicing regulator, and bona fide marker of speckles (Spector & Lamond, 2011). Images of SC35 in uninfected cells showed a large number (typically ~10-15) large nuclear bodies (typical size ~1-2 μm), as expected for speckles (Lamond & Spector, 2003) (**Fig. 3C**). In HIV-1 infected cells, SC35 displayed a similar pattern. Interestingly, nuclear vDNA clusters colocalized with a subset of these SC35 positive bodies (median Pearson’s r=0.14; Costes p<0.01 in 16/17 regions) (**Fig. 3C,D; Fig. S11**). Thus, our data suggests that HIV-1 genome clusters enriched in the paraspeckle protein CPSF6 associate with the splicing and speckle factor SC35.

## Viral integration is not required for viral DNA cluster formation

To determine whether the formation of vDNA clusters requires the integration of the viral genome into host chromosomes, we infected cells with an integration-deficient mutant virus (D116A)(Berger et al, 2009). Interestingly, we again observed the presence of large vDNA clusters which are also associated with vRNA (**Fig. 4A, Fig. S12**). As for the virus containing the functional integrase (**Fig. 2A-C**), the colocalization of vDNA and vRNA was highly significant and intensities in colocalizing clusters correlated significantly (Spearman’s r=0.51, p=1.6×10^-5^) (**Fig. 4B,C**). Additionally, when we treated cells at the time of infection with 10 μM of Raltegravir (RAL), an inhibitor of HIV integration, we also observed large and bright vDNA clusters associated with vRNA (**Fig. S13B**). These experiments demonstrate that the formation of vDNA/vRNA clusters does not depend on integration of the viral genome into host chromosomes.

**Figure 4:**
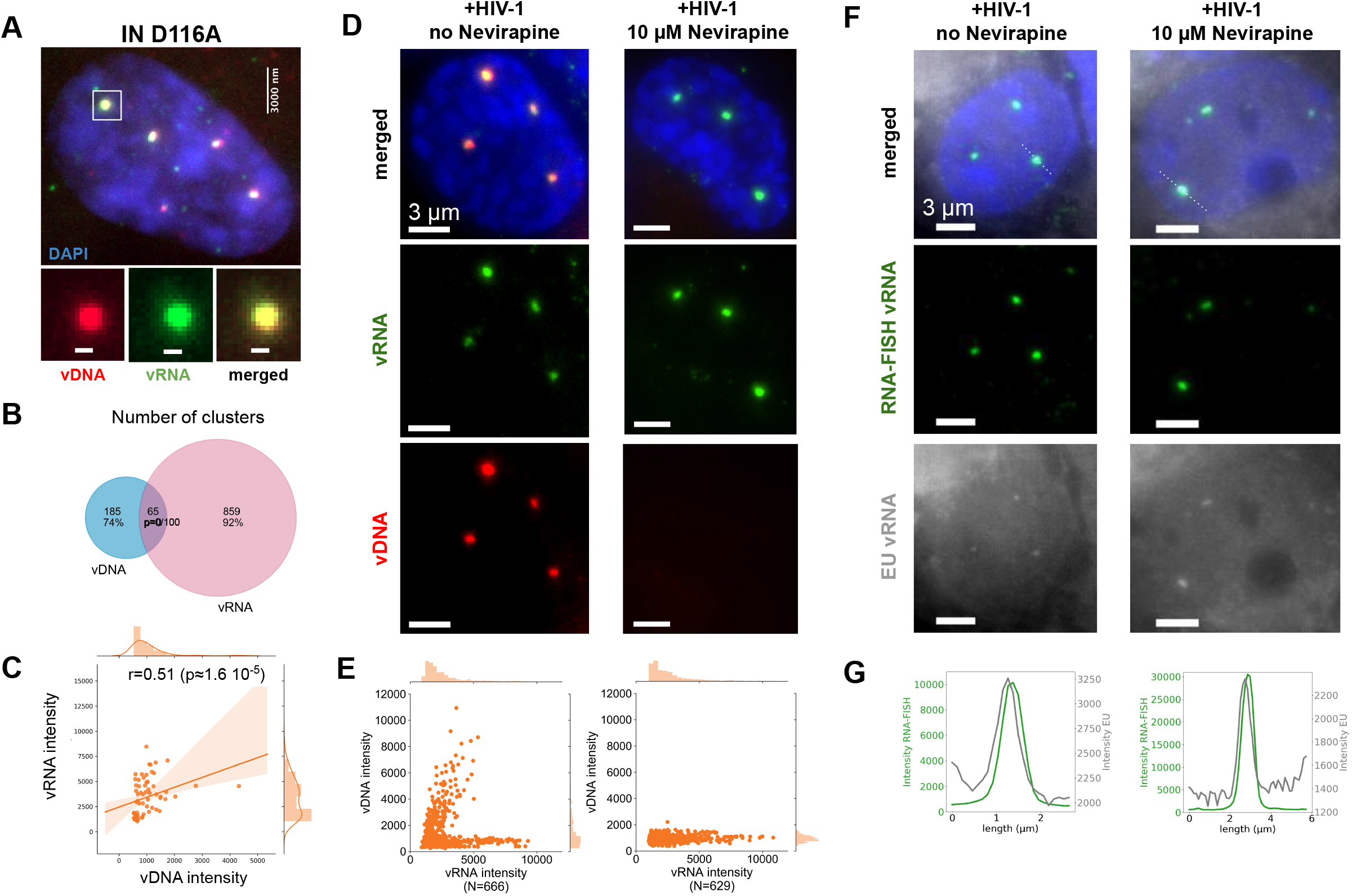
Nuclear clusters can form in absence of integration and contain incoming vRNA. (**A**) Image of a ThP1 cell infected with an integration-deficient HIV-1 carrying the mutation D116A in the catalytic site of IN. The EdU-labeled vDNA is shown in red and the vRNA detected by RNA-FISH in green. The nucleus (DAPI) is shown in blue. See **Fig. S12** for a larger image region. (**B**) Venn diagram shows the number of vRNA clusters, the number of vDNA clusters, and the number of vRNA clusters colocalizing with vDNA clusters. The p-value indicates the significance of colocalization based on a jittering analysis (see Methods). (**C**) Scatter plot shows vDNA and vRNA intensities in colocalizing vDNA and vRNA clusters with the Spearman correlation *r* and associated p-value. (**D**) ThP1 cells were infected with HIV-1 at an MOI of 20, in absence (left) or presence (right) of nevirapine (NVP, 10 μM). Blue: DAPI staining. Green: vRNA (RNA-FISH). Red: vDNA (EdU). (**E**) Scatter plots show intensities of vDNA and vRNA in detected vRNA clusters. (**F**) Image of a ThP1 cell infected by HIV-1 with EU labeled RNA in absence (left) or presence (right) of nevirapine (NVP, 10 μM). Blue: DAPI staining. Green: vRNA (RNA-FISH). Grey: EU-labeled vRNA. See **Fig. S19** for a larger image region. (**G**) Intensity profiles of EU and RNA-FISH along the dotted lines in panel **F** (top).

## Viral RNA clusters contain mostly genomic RNA

The dual color EdU/RNA-FISH images reported above clearly indicate that vDNA clusters colocalize with vRNA (**Fig. 2A-C**). However, our RNA-FISH probes against the HIV-1 POL gene cannot distinguish between incoming (genomic) vRNA and newly transcribed (messenger) vRNA. We therefore sought to verify the nature of the observed vRNA by treating cells with 10 μM of Nevirapine (NVP), a potent RT inhibitor. The efficacy of the drug was demonstrated by the absence of detectable vDNA in infected nuclei; to our surprise, however, we still observed bright vRNA clusters in the nucleus, whether we exposed the cells to NVP for 24 h, 48 h or 72 h (**Fig. 4D,E; Fig. S1, S14A, S20**). Because HIV-1 transcripts cannot be produced in the absence of vDNA, we conclude that nuclear vRNA clusters at these time points contain mostly incoming, genomic vRNA, in NVP treated cells.

We next asked if genomic vRNA clusters also exist in absence of RT inhibition. Intensity distributions of vRNA clusters in NVP treated cells at 48 h p.i. were similar to those in untreated cells (**Fig. S14B**), strongly suggesting that the vRNA clusters in the untreated cells are also mostly composed of genomic vRNA. To verify this directly, we turned to a different labeling technique that, unlike RNA-FISH, highlights the genomic, but not the transcribed, vRNA. Specifically, we used 5-ethynyl uridine (EU), another nucleoside analog that incorporates into nascent RNA during transcription and can be fluorescently detected by click-chemistry, as for EdU (Jao & Salic, 2008; Xu et al, 2013). As demonstrated previously (Xu et al, 2013), we used EU to label HIV RNA in 293T producer cells, then we infected NVP treated or untreated ThP1 cells with this virus (MOI 50), and performed fixation at 24 h p.i., followed by click chemistry labeling and fluorescence imaging, in combination with RNA-FISH (**Fig. 4F, Fig. S15**). Despite the presence of a significant background signal, we observed EU spots in the nucleus of both NVP treated and untreated cells that coincided with vRNA clusters (**Fig. 4G, Fig. S15**). These data therefore strongly support the presence of genomic vRNA in the nucleus, regardless of whether cells have been pharmacologically treated.

## Nuclear viral RNA clusters can undergo reverse transcription

The unexpected presence of genomic vRNA clusters in the nucleus (**Fig. 4D,E,F,G**) raises the intriguing question as to whether they can serve as templates for RT. To test this possibility, we took advantage of the reversibility of RT inhibition by NVP. We reasoned that if RT occurs locally in vRNA clusters, allowing RT to resume after temporary RT inhibition would lead to the detection of newly synthesized vDNA within these nuclear structures.

We first exposed cells to NVP starting from the time of infection during 24, 48 or 72 hours. Dual color images obtained at all three time points showed an absence of nuclear vDNA signal, as expected, while the vRNA clusters remained clearly visible, as above (**Fig. S14A, Fig. 4D**). Next, we exposed cells to NVP starting from the time of infection for 48 h or 72 h, then removed NVP by wash-out and imaged cells 24 h later. Strikingly, in both experiments, we observed clear vDNA clusters that colocalized with the vRNA clusters, and displayed positive correlation of their intensities (**Fig. 5A,B,E,F; Fig. S16**), as in untreated cells (**Fig. 2A-C**). This appearance of vDNA at vRNA clusters can be explained by a local RT activity within the nuclear vRNA clusters. Alternatively, the observed vDNA clusters might reflect nuclear import of vDNA synthesized exclusively in the cytoplasm, between the time points of NVP wash-out and fixation. In the first case (nuclear RT), we expect that clusters containing more vRNA also exhibit higher amounts of vDNA. This is indeed the case, as evidenced by the positive correlations between vRNA and vDNA intensities in colocalizing clusters in all wash-out experiments (**Fig. 5E**).

**Figure 5:**
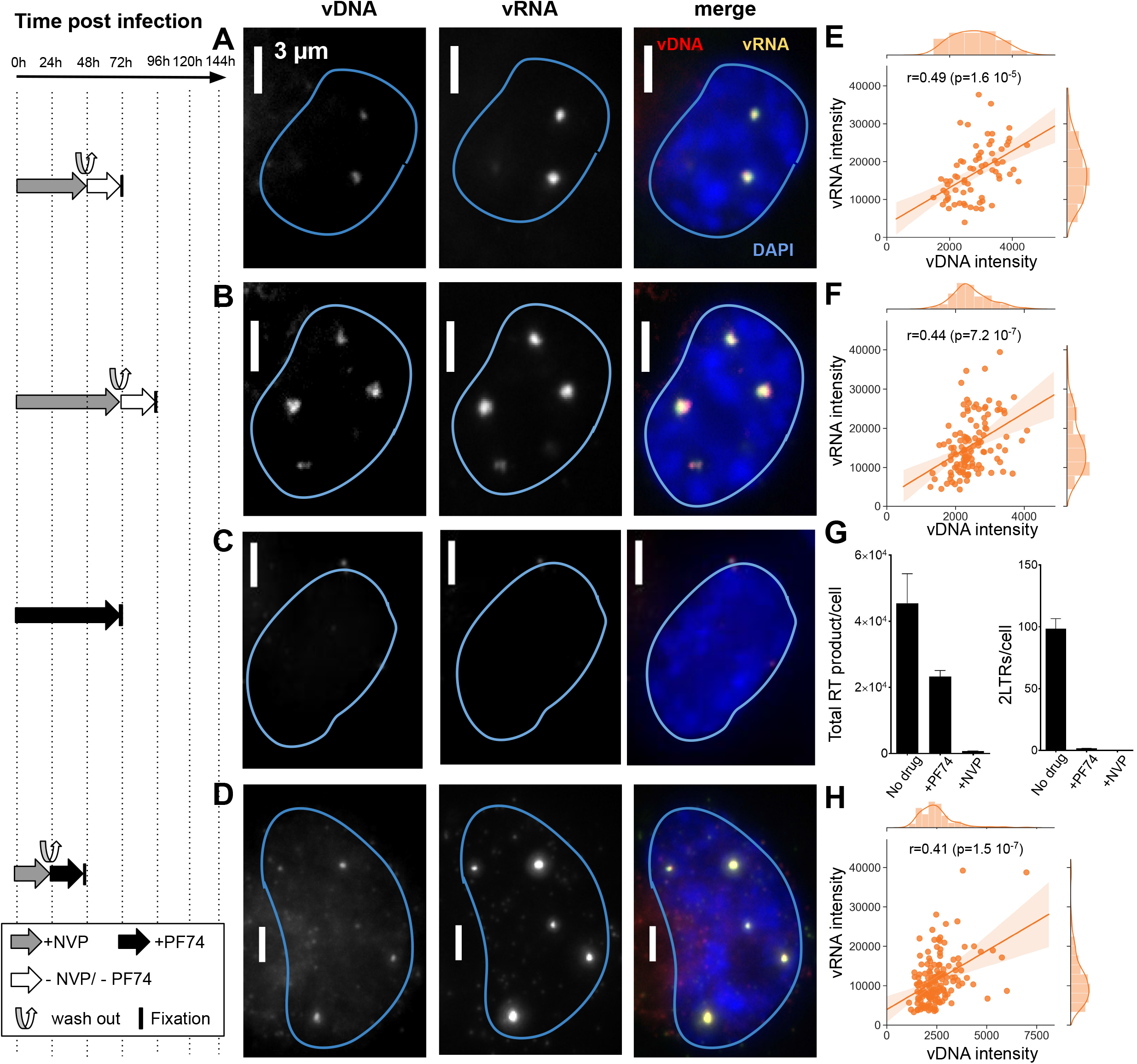
Reverse transcription in nuclear clusters of infected macrophages. The left panel shows the timeline of drug exposure experiments. (**A,B**) ThP1 cells were exposed to NVP for 48 h (**A**) or 72 h after infection (**B**), then NVP was washed out, and cells were fixed for click chemistry 24 h later. (**C**) Cells were exposed to PF74 for 72 h after infection, then fixed for click chemistry. (**D**) Cells were exposed to NVP for 24 h after infection, then NVP was washed out, and cells were exposed to PF74, before being fixed for click chemistry 24 h later. (**A-D**) Images show vRNA and vDNA in an infected ThP1 cell (MOI 20) for each of the four experimental conditions. See **Fig. S15-S17** for images of larger regions. (**E,F,H**) Scatter plots show vDNA and vRNA intensities in colocalizing clusters with Spearman correlation *r* and associated p-values. Replicates of the four experiments yielded similar results. (**G**) qPCR measures DNA synthesis (left) and nuclear import (right) at 72 h p.i. in absence of drug treatment, or after exposure to NVP or PF74.

To further assess both possibilities, we next quantified the amount of vRNA in the cytoplasmic and nuclear compartments using an automated analysis based on FISHquant and ImJoy (Mueller et al, 2013; Ouyang et al, 2019). These analyses indicated that the nuclear vRNA pool exceeded the cytoplasmic pool by at least ~7-fold after 72 h of NVP treatment (**Fig. S14C**). Thus, the amount of vRNA available for cytoplasmic RT and subsequent nuclear import during the 24 h after wash-out is much smaller than the amount of vRNA available for nuclear RT. Therefore, the nuclear vDNA detected in our NVP wash-out experiments more likely arises from local RT in nuclear clusters rather than import of vDNA synthesized in the cytoplasm.

To directly rule out a contribution of cytoplasmically synthesized vDNA to nuclear vDNA clusters after NVP removal, we aimed to pharmacologically block nuclear import using the molecule PF-3450074 (PF74). At low concentrations (1.25-2.5 μM), PF74 impedes nuclear import of HIV-1 without abolishing RT (Blanco-Rodriguez et al, 2020; Bejarano et al, 2019; Francis & Melikyan, 2018; Balasubramaniam et al, 2019; Blair et al, 2010). Indeed, when we exposed cells to 1.5 μM of PF74 for 72 h after infection, qPCR indicated that RT was roughly halved, while 2LTR formation was virtually eliminated (**Fig. 5G**). We observed no discernable nuclear vRNA in PF74 treated cells at 48 h or 72 h p.i. and a higher cytoplasmic vRNA pool as compared to NVP treated cells at the same time point, confirming inhibition of nuclear import (**Fig. 5C, Fig. S17A,B; Fig. S18**). Finally, we treated cells with NVP for 24 h p.i., then washed NVP out and immediately exposed the cells to 1.5 μM PF74 for another 24 h. Despite the block of nuclear import, we again observed clear vDNA clusters colocalizing with vRNA clusters in the nucleus, with positively correlating intensities (**Fig. 5H; Fig. S17C**). We made the same observation when exposing cells to NVP for 48 h, applying PF74 at 36 h, and washing out NVP (while keeping the cells exposed to 1.5 μM of PF74) at 48 h p.i. (**Fig. S19**). We additionally performed qPCR analyses of circular 2LTRs, an exclusively nuclear form of vDNA that can persist in the nucleus for several weeks (Gillim-Ross et al, 2005) (**Fig. S1**). Interestingly, we were able to amplify 2LTRs in samples after NVP wash-out, whether or not followed by exposure to PF74, albeit with a 3-7 fold reduction compared to untreated cells; this reduction was ~10-fold when applying PF74 at 36 h p.i. and washing out NVP at 48 h p.i. (**Fig. S20**). Nevertheless, the detection of 2LTRs in these experiments corroborates the synthesis of complete vDNA by RT in the nucleus. We also analyzed the presence of proviral DNA using ALU-PCR (Lelek et al, 2015; Di Nunzio et al, 2013). While we could amplify these integrated vDNA forms in untreated cells or in simple NVP wash-out experiments, we failed to do so in the experiments involving PF74 exposure (**Fig. S20**). This suggests that the majority of the vDNA detected consist of unintegrated genomes, in agreement with our above findings using an integration deficient virus (**Fig. 4A-C**) or when inhibiting integration pharmacologically (**Fig. S12**). Notwithstanding, our data argue against the possibility that the nuclear vDNA clusters arise exclusively from import of cytoplasmically synthesized vDNA and therefore constitute compelling evidence for a nuclear RT activity within vRNA clusters after the release of RT inhibition.

## Nuclear RT activity results in transcription competent vDNA

In addition, we asked whether the viral DNA synthesized in the nucleus after pharmacological block followed by release of RT is competent for transcription. To address this, we infected cells with the virus carrying the GFP reporter gene used above (**Fig. 2D, S21**) and quantified the percentage of GFP positive cells using flow cytometry (FACS) (**Fig. 6A-D, S21G,H**). Cells infected with MOI 20 and fixed at 3 (**Fig. S21G**) days or 7 days (**Fig. 6B**) p.i. displayed ~ 9% and ~12% of GFP positive cells, respectively, vs. 0% for uninfected cells (**Fig. 6A**). When we inhibited RT by NVP treatment for 3 days, allowed RT to resume by washing out NVP and fixed cells for imaging 4 days later, ~4% of cells were found to be GFP positive (**Fig. 6C**) against 0.3% of GFP+ cells in presence of NVP at 7 days p.i. (**Fig. S21H**). The percentage of GFP positive cells was lower when 1.5 μM of PF74 was added at the moment of NVP wash-out, or 12 h before the wash-out, to block nuclear import of the remaining cytoplasmic virus (~1.2% and ~0.94%, respectively) (**Fig. 6D, Fig. S21H**). Although these percentages are considerably lower than for untreated cells, they are much larger than for cells treated with NVP until fixation (0.04% at 3 days p.i., 0.33% at 7 days p.i.) (**Fig. S21G,H**) or cells treated with PF74 for 3 days (0.15%) (**Fig. S21G**). These FACS results were corroborated by imaging (**Fig. 6E-H, Fig. S21A**). Importantly, we did not use EdU in these experiments because we observed that EdU negatively interferes with viral transcription (**Fig. S21F**). Overall, our data suggest that nuclear RT activity, at least in conditions where RT was initially suppressed, can lead to the production of transcription competent viral DNA.

**Figure 6:**
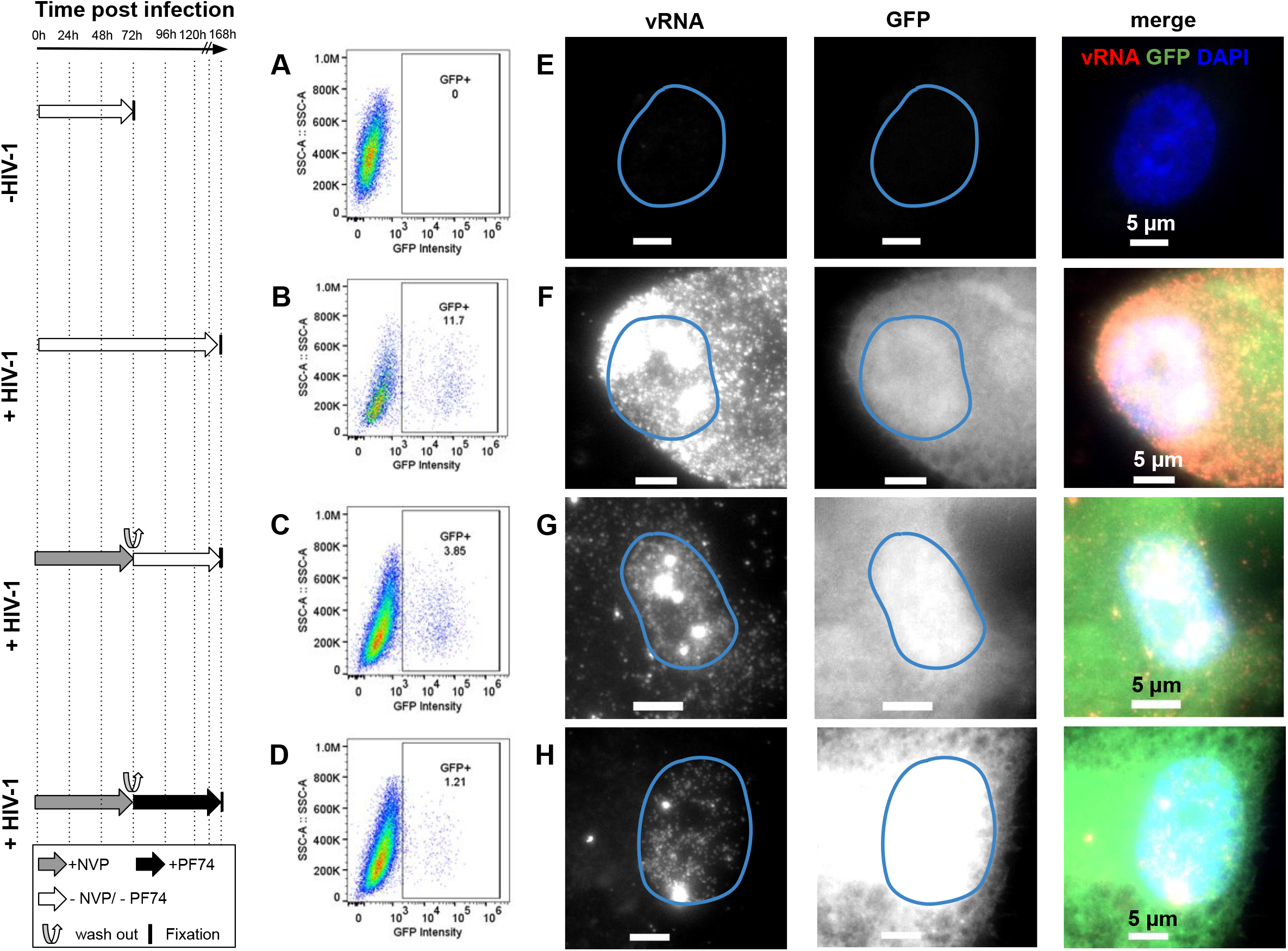
Nuclear RT can yield transcription competent vDNA. (**A-D**) Percentage of GFP positive cells at 3 and 7 days post-infection analyzed by FACS in absence of EdU. (**E-H**) Multicolor images of ThP1 cells infected with a GFP-reporter virus. vRNA (RNA-FISH) is labeled in red, GFP in green, and nuclei (DAPI) in blue. (**A,E**) Uninfected control cells. (**B,F**) Untreated, infected ThP1 cells fixed at 3 d (72 h) p.i. (**C,G**) Infected cells were exposed to NVP for 72 h, then NVP was washed out and cells were cultured for another 96 h and fixed at 7 d p.i. (**D,H**) Infected cells were cultured for 72 h p.i., then NVP was washed out and cells were exposed to the nuclear import inhibitor PF74 for another 96 h and fixed at 7 d p.i.

## Nuclear clustering and reverse transcription in primary macrophages

Finally, we asked if the main phenotypes described in ThP1 cells above also extend to primary macrophages. We therefore prepared monocyte-derived macrophages (MDM) obtained from two healthy donors. FACS analysis indicated that ~76% of these cells were CD14 positive, confirming that the large majority consisted of macrophages (**Figure S22**). We then infected these MDMs with VSV-G-pseudotyped HIV-1ΔEnv. Cells from donor 1 were infected with the GFP reporter virus in presence of Vpx, while cells from donor 2 were infected with a virus without GFP and in absence of Vpx. We imaged the vDNA and vRNA using EdU and RNA-FISH, using similar experimental conditions as for the ThP1 cells above, including absence of treatment, treatment with NVP, and treatment with NVP followed by wash-out, (**Figure 7** and **Figures S23,EV4**). In untreated MDMs at 7 d post-infection, we again observed bright nuclear foci of vDNA in a sizeable fraction of the cell population for both donors (15 out of 31 counted for cells, i.e. ~50%, for cells, for donor 2 and 22 out of 53, i.e. ~30% for donor 1), thus both in presence and in absence of Vpx (**Figure 7A,B** and **Figure S23A,E**). Focusing on cells from donor 2 (infected without Vpx), we measured vDNA foci sizes (FHWM) similar to those measured in ThP1 cells (median ~520 nm, interquartile range=259 nm, n=20 regions of interest) (**Figure 7E**). These vDNA foci also partially colocalize with vRNA foci (**Figure 7A,B**), as previously shown for ThP1 cells (Costes p-values <0.05 in 18 out of n=20 regions) (**Figure 7F**). Neither vDNA nor vRNA foci could be observed in uninfected MDM cells (**Figure S23B,F**).

**Figure 7:**
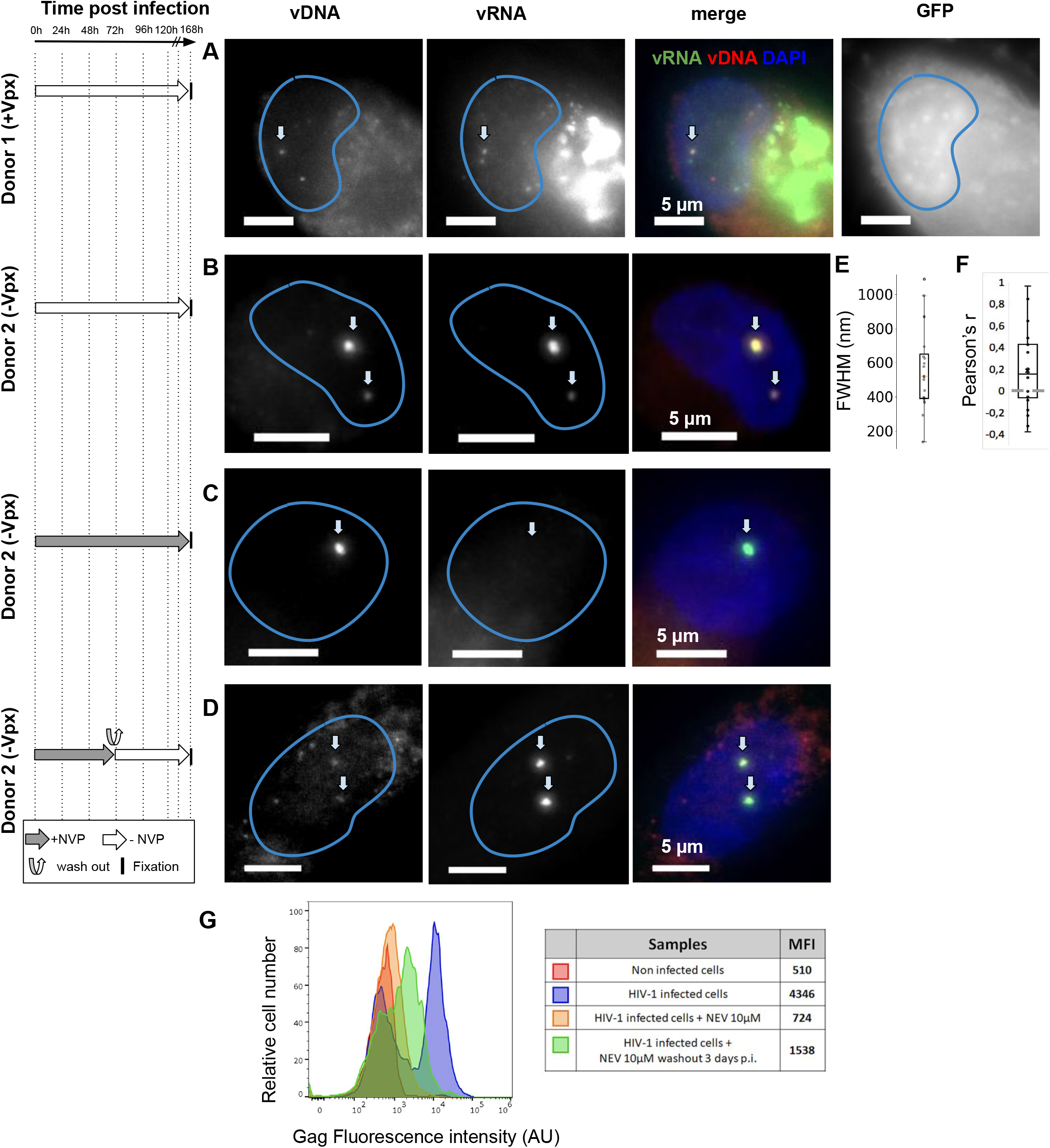
Clustering of vDNA/vRNA and nuclear RT in primary human macrophages. **(A-D)** Multicolor images of vDNA, vRNA, DAPI and/or GFP in monocyte derived macrophages (MDMs) from two different donors, infected with HIV1. Cells from donor 1 were infected with VSV-G-pseudotyped HIV-1ΔEnv Vpx carrying a GFP reporter (**A**) and cells from donor 2 were infected with a VSV-G-pseudotyped HIV-IΔEnv without Vpx (**B-D**). (**A,B)** Infected cells were left untreated and fixed at 6 d p.i. (**C**) Infected cells were treated with 10 μM Nevirapine (NVP) throughout the experiment and fixed at 6 d p.i. (**D**) Infected cells were exposed to NVP for 3 d p.i., then NVP was washed out and cells were cultured for another 4 d and fixed at 7 d p.i. (**E**) Size of DNA foci in untreated infected cells from donor 2 imaged at 7d p.i. (**B**), measured by the FWHM of n=24 foci. (**F**) Pearson’s r showing colocalization of DNA with RNA foci in the same cells (n=20). **(G)** FACS analysis of Gag positive cells from donor 2 in different conditions. Red: uninfected cells. Blue: untreated infected cells at 7 d p.i. as in **B**. Orange: cells were treated with NVP for 7 d as in **C**. Green: infected cells were exposed to NVP for 3 d, then NVP was washed out and cells were cultured for another 4 d as in **D**. Median of fluorescence intensities (MFI) for each sample are shown in the table.

MDM cells of donor 2 treated with 10 μM of NVP for 7 d also displayed vRNA foci, but not vDNA foci, indicating that genomic vRNA accumulates in nuclear foci of MDMs much as in ThP1 cells (**Figure 7C, Figure S23C**). Moreover, in MDM cells treated with NVP for 3 days, followed by wash-out and fixation 4 days later, we again observed vDNA signal in vRNA clusters, consistent with a resumption of nuclear RT activity as previously observed for ThP1 cells (**Figure 7D, Figure S23D**). Similar observations were made for donor 1 (**Figure EV4**).

To determine if nuclear RT can lead to a transcriptionally competent vDNA template, we used FACS to analyze production of the HIV polyprotein Gag in MDMs from donor 2. Our data indicate that under the conditions used above to determine nuclear RT (NVP treatment for 3 d followed by wash-out and 4 d recovery), Gag median fluorescence intensity (MFI) is larger than in uninfected cells or cells treated with 10 μM NVP for 7 d (1538 vs. 510 and 724, respectively) (**Figure 7G**). Under the same experimental conditions, we also observed some GFP positive cells in MDMs from donor 1 (**Figure EV4**). These data support the notion that nuclear RT in MDMs can lead to synthesis of transcriptionally competent vDNA.

Thus, the main phenotypes detailed above in ThP1 cells can also be observed in MDMs, irrespective of the presence of Vpx. In summary, our data show that in primary macrophages, HIV-1 RNA genomes form RT competent nuclear clusters that can lead to transcriptionally competent viral DNA.

## Discussion

Our study uses imaging of viral DNA and RNA to shed new light on the early replication cycle of HIV-1 in macrophage-like (ThP1) cells and in primary (monocyte-derived) macrophages. We demonstrated that vDNA and vRNA genomes cluster together in nuclear niches associated with speckle factors and that these clusters can form in absence of viral integration into the host genome. We further showed that genomic vRNA clusters can form in absence of vDNA after pharmacological inhibition of RT, but, importantly, also in untreated cells. Our observation of genomic RNA in nuclei agrees with previous studies showing that RT is dispensable for nuclear import (Bejarano et al, 2019; Burdick et al, 2013, 2017, 2020). However, the potential role of genomic vRNA in the nucleus remained unknown. Several recent studies have evoked the possibility of RT in the nucleus (Bejarano et al, 2019; Burdick et al, 2017, 2020). However, our experiments combining reversible RT inhibition with direct visualization of the synthesized vDNA provide the first clear demonstration that these genomic RNA clusters can serve as templates for RT in the nucleus. We additionally showed that the viral DNA resulting from this nuclear RT activity can serve as a template for transcription. We note that two recent studies published during the revision of this paper reported similar findings (Francis et al, 2020; Dharan et al, 2020).

We emphasize that our data do not implicate that RT occurs exclusively, or even majoritarily, in the nucleus. Our results allow the possibility that RT is initiated in the cytoplasm and prolonged in the nuclear compartment, or that some viral RNA genomes are entirely reverse transcribed in the cytoplasm. We also acknowledge that our demonstration of nuclear RT was achieved after temporary pharmacological inhibition of RT. Nevertheless, our observation of nuclear genomic vRNA clusters in absence of drug treatment opens the possibility that RT also occurs in the nuclei of untreated macrophages. At any rate, our clear evidence for nuclear RT under the aforementioned conditions stands in stark contrast with the classical picture of the HIV replication cycle, according to which RT is entirely restricted to the cytoplasmic compartment. An intriguing speculation is that these vRNA clusters may act as nuclear microreactors that concentrate reverse transcriptase enzymes to enable efficient vDNA synthesis in macrophages, much as intact capsid cores are believed to concentrate these enzymes in the cytoplasm of HeLa or CD4+ cells (Warrilow et al, 2007).

Previous *in vitro* experiments have suggested a potential link between HIV-1 and nuclear speckles (Bell et al, 2001; Pendergrast et al, 2002) and earlier studies have reported that HIV-1 RNA colocalizes with the speckle factor SC35 (Cardinale et al, 2007; Bøe et al, 1998). Here, we directly show that HIV-1 RNA/DNA genome clusters localize to nuclear niches enriched in SC35. While it has been proposed that the nuclear pool of unspliced HIV-1 RNAs may be stored in nuclear paraspeckles (Zhang et al, 2013, 1), our data instead indicate that viral clusters sequester the host cell factor CPSF6 away from its canonical localization in paraspeckles (Cardinale et al, 2007; Bøe et al, 1998; Burdick et al, 2020) and relocate this protein in speckles. Indeed, a recent study demonstrates the role of CPSF6-CA interactions in mediating the association of HIV-1 with nuclear speckles (Francis et al, 2020).

Other studies highlighted the ability of other viruses to alter the cytoplasmic or nuclear organization and form microenvironments that locally concentrate viral and host cell factors required for the synthesis of viral progeny and/or to protect the virus from cellular defense mechanisms (Schmid et al, 2014; McSwiggen et al, 2019; Heinz et al, 2018). One intriguing, albeit speculative, possibility is that the crowded microcompartments formed by cellular RNAs in nuclear bodies shroud the viral DNA and protect it from mediators of the innate immune response, such as the cyclic GMP-AMP synthase (cGAS), a viral DNA sensor that associates with HIV DNA in nuclei of infected macrophages (Lahaye et al, 2018; Sumner et al, 2019).

Our observation of vRNA/vDNA clusters in cells infected by an integration-defective virus or upon inhibition of integration further suggests that most of the vDNA in nuclear clusters may be unintegrated. This dovetails with the common observation of large amounts of unintegrated viral DNA in the nucleus, which can act as viral reservoirs and constitute an important obstacle to successful treatment of HIV-1 infection(Bell et al, 2001; Hamid et al, 2017; Gelderblom et al, 2008). Thus, the unintegrated vDNA observed in our study may be relevant to understanding HIV reactivation in patients.

More research is needed to further explore the functional role and formation mechanisms of these vRNA/vDNA clusters. Our discovery of RT-competent clusters of viral RNA and DNA in ThP1 cells and primary macrophages opens new perspectives for understanding and hence combating HIV-1 replication in natural target cells.

## ACKNOWLEDGEMENTS

We thank Mickaël Lelek and Andrey Aristov for help with microscopy and Benoît Lelandais for help with image analysis. We thank Marie-Anne Welti, Edouard Bertrand, Xavier Darzacq, Olivier Schwartz, Arnaud Echard and Michaela Mueller for useful discussions and/or comments on the manuscript. We thank Fabrizio Mammano for kindly providing the modified IN-HA HIV-1 virus and Stella Frabetti for technical support. We also thank the UtechS Photonic BioImaging, C2RT, Institut Pasteur, which is supported in part by the French National Research Research Agency (France BioImaging; ANR-10*INSB*04; Investments for the Future). We thank the NIH AIDS Reagents program for reagents. This work was funded by Institut Pasteur, Fondation pour la Recherche Médicale en France (DEQ 20150331762), Institut Carnot Pasteur MS, FRM/Sidaction (VIH20170718001) and ANRS (grants N. ECTZ4469-ECTZ74440-ECTZ88162).

## Conflict of interest

None declared

## EXPANDED VIEW FIGURE LEGENDS

**Figure EV1:**
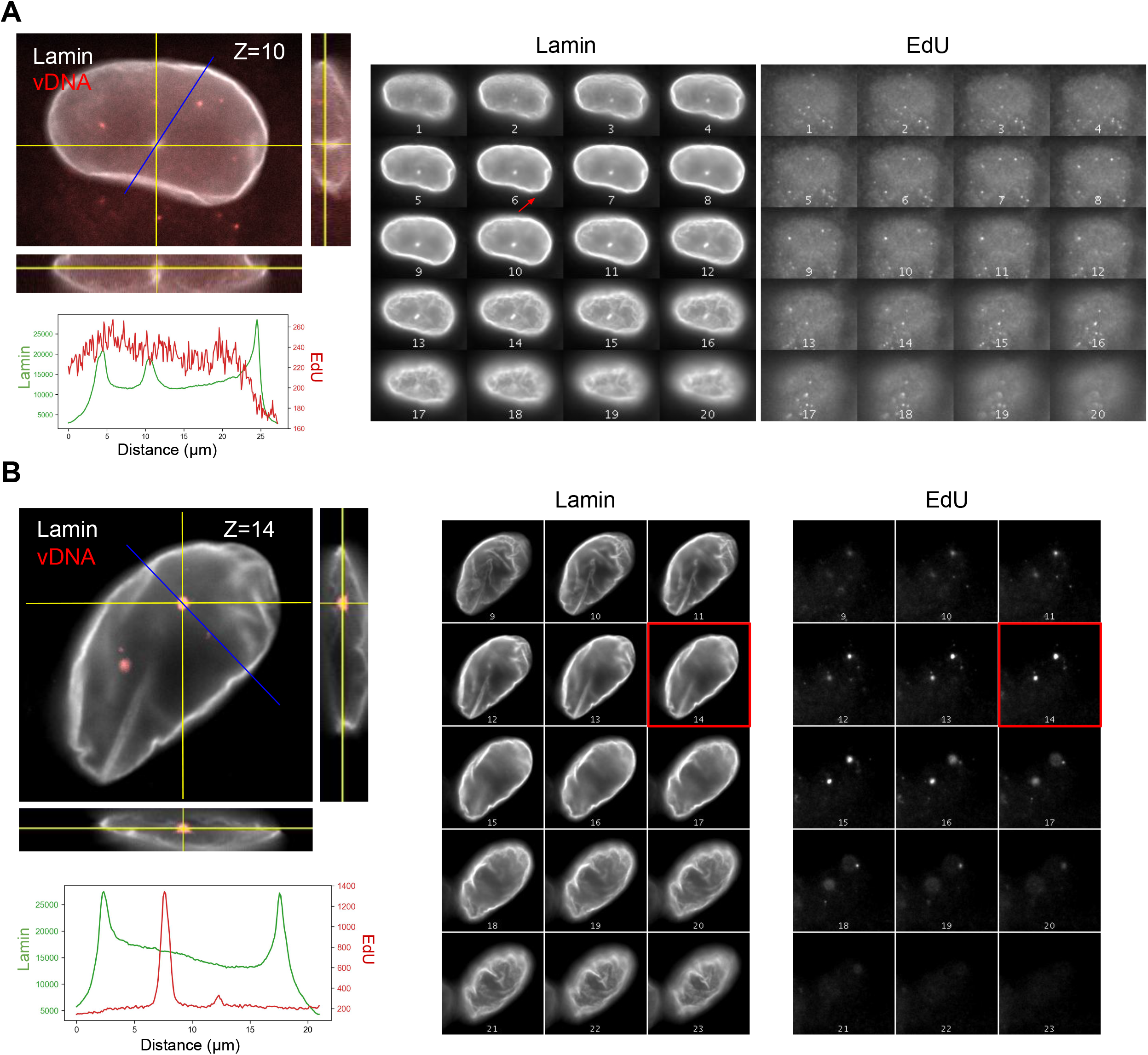
EdU foci are not located in nuclear invaginations. (**A,B**) 3D images of immunostained lamin and EdU are shown to assess the presence of nuclear envelope invaginations and the location of EdU foci in infected ThP1 cells. Images on the left are orthogonal views of z-stacks, where the central image shows an XY slice, and the images to the right and below show XZ and YZ slices at positions indicated by the yellow lines. The blue line indicates the location for the measurement of intensity profiles. The plots below show intensity profiles for the Lamin and EdU channel along these blue lines. Montage views on the right show all z-slices of the Lamin and EdU channels separately. (**A**) Nucleus of an infected cell showing an invagination, as evidenced by the enrichment of Lamin in the intensity profile (middle peak), as well as in the montage views. The chosen line profile crosses the invagination but does not show an enrichment of EdU. (**B**) Another infected cell with a line profile chosen to cross one of the EdU foci. No lamin enrichment was detected at the location of the EdU foci (peak of EdU). Panels on the left are identical to Fig. **1 C**.

**Figure EV2:**
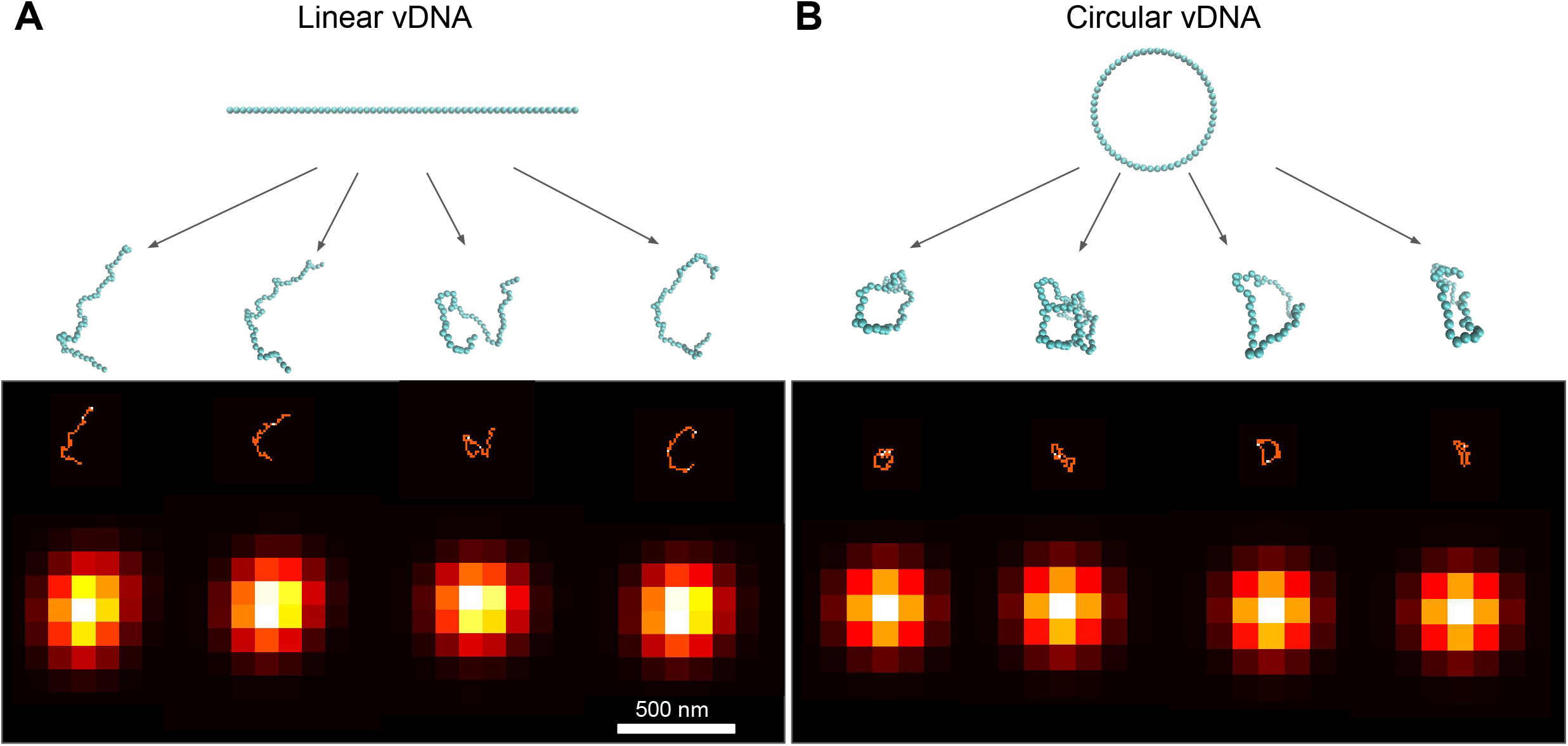
Simulated configurations and images of HIV-1 DNA. (**A,B**) Simulated configurations and images of the HIV-1 DNA in linear (**A**) or circular form (**B**). The 10 Kb long viral genome is represented as a chain of 59 nucleosomes of 11 nm diameter. Starting from the linear or circular initial structures shown (blue chains on top), molecular dynamics simulations (Langevin dynamics) generate 100 independent configurations (only 4 are shown as blue chains of beads). The corresponding image is blurred by convolution with the microscope point spread function (approximated as a Gaussian of standard deviation 100 nm), resulting in the images shown in the bottom row.

**Figure EV3:**
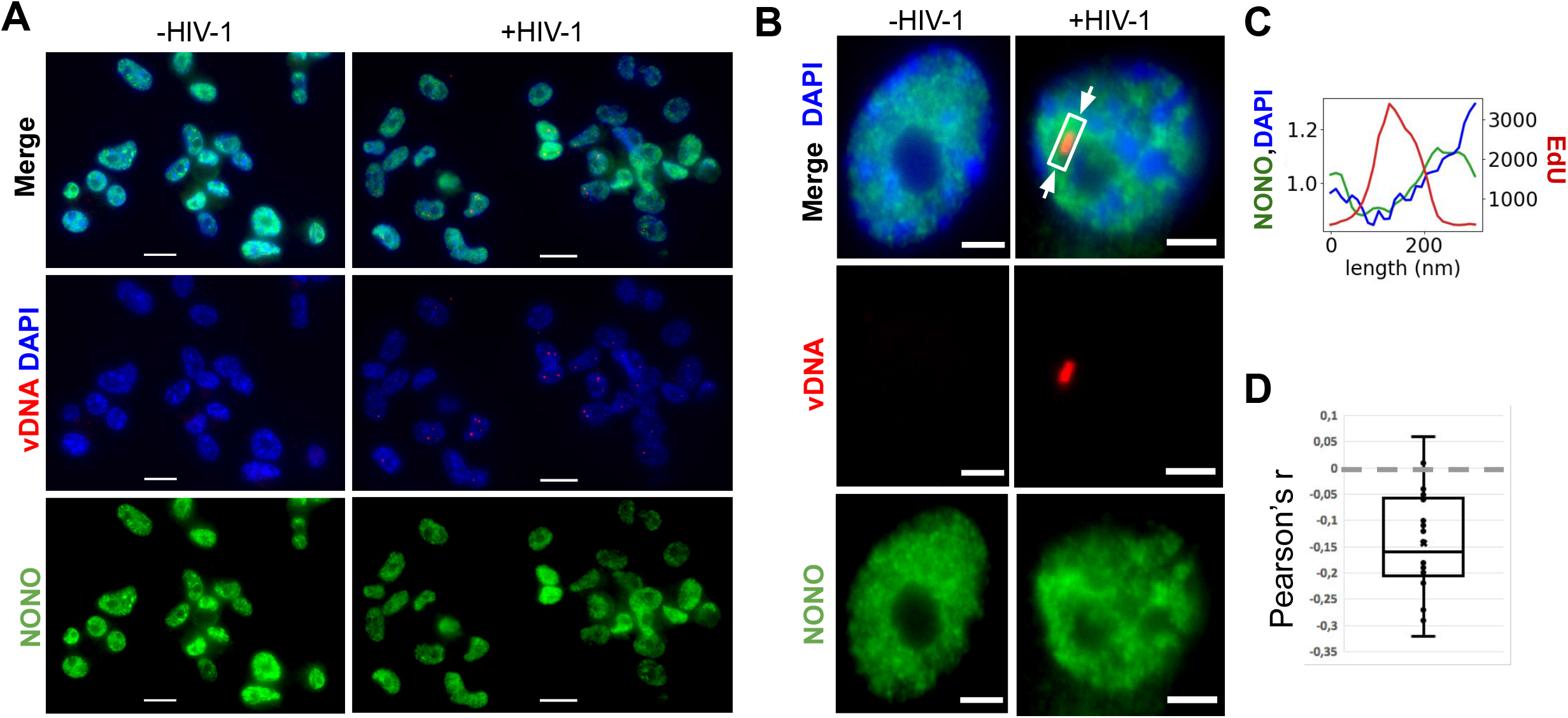
vDNA clusters do not colocalize with the CA-binding protein NONO. (**A,B**) Image of uninfected (left) and infected (right) ThP1 cells showing immunolabeled NONO in green, EdU in red, and the nucleus (DAPI) in blue. (**C**) Intensity profiles of EdU, DAPI and NONO along a profile crossing a vDNA focus (rectangle in B). (**D**) Boxplot shows Pearson correlations between vDNA (EdU) and NONO. The negative correlation values indicate an absence of colocalization between vDNA and NONO (Costes p-value<0.05 for 18 out of 18 cells, and <0.01 for 14 out of 18 cells).

**Figure EV4:**
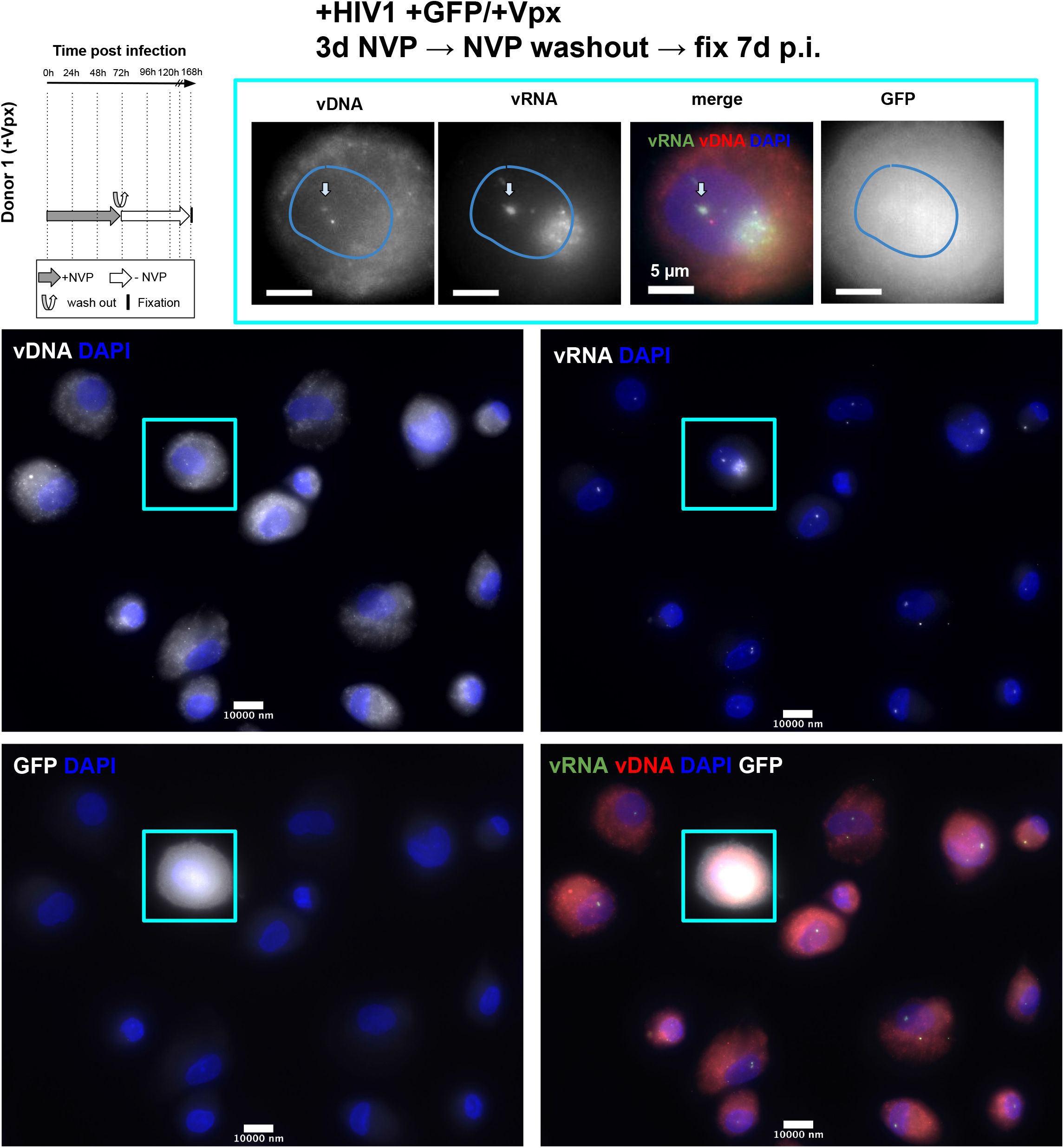
HIV-1 transcription in primary macrophages after temporary inhibition of RT. Multicolor images of MDMs from donor 1, infected with a HIV-1 virus carrying a GFP reporter and in presence of Vpx. Cells were treated with Nevirapine (NVP) for 3d, then NVP was washed out and cells were fixed for imaging 4d later (i.e. at 7 d p.i.). Images show the vDNA, vRNA and/or GFP signal and DAPI.

## METHODS

### Cell culture

Human ThP-1 cells (ATCC TIB-202) were grown in RPMI 1640 medium supplemented with 10 % (vol/vol) fetal bovine serum and 1% (vol/vol) penicillinstreptomycin. For infections, 12-well plates containing coverslips were seeded with 1×10^6^ THP-1 cells and treated with PMA (259 μg/ml final concentration) at 37 °C and 5% CO_2_. Twenty four hours post-stimulation, non-adherent cells were removed, adherent cells were washed, and further cultured. PMA was present during the entire experiment.

### Plasmids and Viral production

The plasmid HIV-1ΔEnv IN_HA_ (D116A)ΔNef was obtained by insertional mutagenesis using the QuikChange II XL Site-Directed Mutagenesis kit (Agilent). HIV-1 viruses were produced by cotransfection with calcium phosphate with 10 μg HIV-1 LAI (BRU) (or NL4.3) ΔEnv Virus (NIH) or with the modified versions HIV-1ΔEnvIN_HA_ (Petit et al, 1999) or HIV-1ΔEnv IN_HA_ (D116A)ΔNef in combination with 1 μg of VSV-G envelope expression plasmid pHCMV-G (VSV-G) and 3 μg of SIV_MAC_ Vpx (Durand et al, 2013). The viruses collected from 293T cells 48 h post-transfection were ultracentrifuged at 4 °C for 1h at 22,000 rpm. Virus normalizations were performed by p24 ELISA according to the manufacturer’s instructions (Perkin Elmer) or by qPCR. HIV-1 ΔEnv Δnef LUC and HIV-1 ΔEnv Δnef GFP viruses were produced and titered similarly.

### Quantitative PCR

DNA synthesis, nuclear import and integration during HIV-1 infection in ThP1 cells were quantified by qPCR. We analyzed for late reverse transcription (LRT) products representing HIV-1 DNA synthesis in the cell and 2LTRs to measure nuclear import by qPCR. Integration of proviruses into the human genome were measured by ALU PCR. Viruses were treated for 30 min at 37°C with 1,000 U of DNase I (Roche). As a control, 10 μM nevirapine was used in infected cells. Total cellular DNA was isolated using the QIAamp DNA micro kit (Qiagen) at 7 and 24 h p.i. Viral DNA synthesis products and 2LTRs were measured at different time points by real-time PCR. LRT, 2LTRs and Alu PCR were performed as described (Lelek et al, 2015; Di Nunzio et al, 2013). LRT were amplified using the primers MH531 and MH532, with the standard curve prepared using the plasmid coding for the viral genome. 2LTRs were amplified by primers MH535/536 and probe MH603, using as standard curve the pUC2LTR plasmid which contains the HIV-1 2LTR junction. Integration was assessed by Alu-PCR, using primers designed in the U3 region of LTR (Lelek et al, 2015; Di Nunzio et al, 2013). The standard curve was prepared as follows: DNA generated from infected cells was end point diluted in DNA prepared from uninfected cells and serial dilutions were made. The control of the first round PCR was the amplification without Alu primers but only U3 primers (Lelek et al, 2015; Di Nunzio et al, 2013). Dilutions of the first round were processed by real time PCR (Lelek et al, 2015; Di Nunzio et al, 2013). All experiments were carried out using internal controls such as infection in presence of RAL (10μM) and/or NVP (10μM). LRT, 2-LTR and Alu-PCR reactions were normalized by amplification of the Actin gene (Lelek et al, 2015; Di Nunzio et al, 2013).

### Virus infection

Cells were infected at least 24 h after initial PMA stimulation with MOIs 10, 50 or 100, based on the viral titer calculated on 293T cells (8.93×10^8^ TU/mL) by qPCR. For experiments with antiretroviral drugs, cells were infected with HIV-1 in the presence of 10 μM RAL or 10 μM NVP or 1.5 μM PF74.

### Drug washout experiment

Cells were grown and differentiated as described above. Cells were infected with an MOI of 50 in the presence of EdU and NVP or PF74. For experiments where NVP was washed out, the drug-containing medium was removed, cells were washed twice with 1 mL of medium for 15 min, fresh medium was added and cells were incubated for the indicated amount of time. The medium was changed every 24 h and reconstituted with or without the PF74 drug, as indicated.

### Labeling of viral DNA by click chemistry and immunolabeling

For imaging of vDNA, CA and IN, ThP1 cells were seeded in 12-well plates on cover glass in the presence of PMA and infected on the following day with HIV-1 at different MOI in medium containing 5 μM EdU. After 24 hours, the medium was removed and replaced by fresh medium containing 5 μM EdU and incubation was continued at 37 °C. To stop the infection, cells were washed with warm PBS and fixed with 4% paraformaldehyde in PBS for 20 min at room temperature.

For vDNA labeling, cells were washed twice with PBS supplemented with 3% bovine serum albumin (BSA) and permeabilized with 0.5% (vol/vol) Triton X-100 for 30 min. After washing with 3% BSA in PBS twice, click-labeling was performed for 30 min at room temperature using the Click-iT EdU-Alexa Fluor 647 Imaging Kit (Thermo Fisher Scientific) following the manufacturer’s instructions.

For immunolabeling, cells were blocked for 30 min with 3% BSA in PBS and permeabilized with 0.5% (vol/vol) Triton X-100 for 30 min. After 2 washes with 3% BSA in PBS, cells were incubated with the primary antibody in 1% BSA in PBS for 1 h at room temperature. After washing with 1% BSA in PBS, a secondary antibody was used for 1 h at room temperature in 1% BSA in PBS. The primary and secondary antibodies used in this study are listed in **Note S1**.

In experiments that combine EdU and protein labeling, the click chemistry reaction was performed prior to immunolabeling and blocking with 3% BSA was omitted.

### Labeling of viral RNA with single molecule RNA-FISH

To visualize individual vRNA molecules, we used the smiFISH approach (Tsanov et al, 2016). Unlabelled primary probes are designed to target the RNA of interest, and can be pre-hybridized with fluorescently labeled secondary detector oligonucleotides for visualization. All probes, except against Neat1, were designed using either OLIGOSTAN (Tsanov et al, 2016) or Stellaris Probe designer, and purchased from Integrated DNA Technologies (IDT). All probe sequences are available in **Tables S1-3**. We used 24 primary probes, each 18-20 nt long, against the HIV-1 POL gene. Probe sets against GFP and LUC comprised 18 and 22 probes, respectively. Secondary probes are conjugated to either Cy3 or Cy5. To detect Neat1, we used Stellaris FISH probes against human NEAT1 5’ segment with QUASAR 570 DYE SMF-2036-1.

Cells were fixed as described above, washed twice with PBS and stored in nuclease-free 70% ethanol at −20 °C until labeling. On the day of the labeling, the samples were brought to room temperature, washed twice with wash buffer A (2x SSC in nuclease-free water) for 5 min, followed by two washing steps with washing buffer B (2X SSC and 10% formamide in nuclease-free water) for 5 min.

The target-specific primary probes were pre-hybridized with the fluorescently labeled secondary probes via a complementary binding readout sequence. The stock concentration of the probes were: Luciferase LUC: 148.6 ng/μL, POL: 1497 ng/μL and GFP: 163.3 ng/μL.

The reaction mixture contained primary probes at a final concentration of 40 pm, and secondary probes at a final concentration of 50 pm in 1x NEB3 buffer. Prehybridization was performed in a PCR machine with the following cycles: 85 °C for 3 min, followed by heating to 65 °C for 3 min, and a further 5 min heating at 25 °C. 2 μL of this FISH-probe stock solution was added to 100 μL of hybridization buffer (10% (w/v) dextran, 10% formamide, 2X SSC in nuclease-free water).

Samples were placed on Parafilm in a humidified chamber on 100 μL of hybridization solution, sealed with Parafilm, and incubated overnight at 37°C. The next day, cells were washed in the dark at 37°C without shaking for >30min twice with pre-warmed washing buffer B. Sample were washed once with PBS for 5 min, stained with DAPI in PBS (1:10000) for 5 min, and washed again in PBS for 5 min. Samples were mounted in ProLong Gold antifade mounting medium.

For simultaneous detection of viral DNA with click chemistry and vRNA with RNA-FISH, the click reaction was performed prior to RNA-FISH and as described above, except no BSA was used in order to minimize RNA degradation by RNases. The ratio copper-protectant/copper solution was 1.5:1 to avoid degradation of RNA, as indicated by the manufacturer.

### Monocyte isolation, differentiation and infection

Peripheral blood mononuclear cells (PBMCs) were isolated from two HIV-seronegative donors (Etablissement français du sang, EFS), by density-gradient centrifugation. Monocyte derived macrophages (MDMs) were prepared by adherence with washing of non-adherent cells after 2 h. Adherent cells were selected in RPMI 1640 medium supplemented with 10% human serum or 10% foetal calf serum (FCS) and MCSF (10 ng/ml) for 3 days and then differentiated for another 4 days in RPMI 1640 medium supplemented with 10% FCS without MCSF. FCS was used to prepare stimulated cells following the protocol described in Mlcochova et al. (Mlcochova et al, 2017). MDMs of donor 1 were infected with MOI 100 of HIV-1 GFP in the presence of Vpx. MDMs of donor 2 were infected with 500ng of p24 of the HIV-1 without Vpx. Samples in which RT was inhibited, 10 μM NVP was present for indicated periods of time. Samples used for click chemistry staining contained 5 μM final concentration of EdU. MDMs were fixed in PFA 4% for 20 minutes at RT, then washed twice with PBS. Samples were frozen at −20 degrees in 70% cold ethanol for at least 12h before click chemistry and RNA-FISH (see sections above). Duplicates of samples without EdU were analysed by FACS and stained in permeabilized samples with 0.05% Saponin using anti-Gag antibody conjugated to Rhodamine (Beckman Coulter, #6604667) diluted 1:500 in PBS-BSA 0.5%. MDMs were controlled for the expression of CD14 (Ab anti CD14-FITC, BD #555397, dilution 1:20) in unfixed cells.

### Polymer simulations

To estimate the expected size and images of single vDNA particles, we performed polymer simulations in which the vDNA molecule is represented by a semiflexible chain of beads undergoing random motions (Langevin dynamics) (Arbona et al, 2017). We assumed chromatinized (i.e. nucleosome-containing) DNA and performed two types of simulations: one with a linear chain (as expected e.g. for integrated vDNA) and the other with a circular chain (e.g. for unintegrated, episomal vDNA). Each bead represents a single nucleosome consisting of 175 bp of DNA (nucleosomal + linker DNA) confined in a 11 nm spherical bead. We assumed the bending stiffness of the polymer to be 11 nm (single bead) and connected consecutive beads via spring potentials. We started our simulations from a linear or circular configuration and allowed the polymer to reach equilibrium (as judged by the temporal evolution of the gyration radius) before sampling one configuration for each simulation. Simulations were run using the LAMMPS simulation library (https://lammps.sandia.gov/). Simulated images were obtained by convolving the simulated configuration with a 3D Gaussian kernel of standard deviation 100 nm approximating the microscope point spread function. We used 100 independent simulations to compute the distributions of sizes (FWHM) for each model.

### Imaging and analysis

Three-dimensional image stacks with a z-spacing of 200 nm were captured on a wide-field microscope (Nikon TiE Eclipse) equipped with a 60X 1.4 NA objective and an sCMOS camera (Hamamatsu Orcaflash 4), with a lumencor SOLA LED light source with adequate Semrock filters, and controlled with MicroManager 1.4.

Nuclei were automatically detected by a trained neural network implemented within the computational platform ImJoy (Ouyang et al, 2019) (Plugin DPNUnet). Spots in Edu and FISH images were detected with a standard spot detection approach: images were first filtered with a Laplacian of Gaussian filter (LoG), then spots were identified with a local maximum filter with a user defined minimum intensity.

Colocalization analysis between two channels was implemented in a custom ImJoy plugin. Only spots located within the detected nuclei were considered. Colocalization was implemented as a linear assignment problem (Python function linear_sum_assignment from SciPy), where each spot from the first channel is assigned to one spot in the second channel. Each spot can be only assigned once, and only assignments below a user defined distance threshold are permitted. Spot intensities for co-localized spots were measured as the maximum intensity in a +/- 1 pixel window. To calculate p-values for the reported co-localization, the colocalization analysis was repeated 100 times after randomly moving (“jitter”) detected spots in one channel by 1000 nm in XYZ. The reported p-values are the proportion of jitter iterations where a higher colocalization was obtained than in absence of jitter.

EdU enrichment for Nevirapine experiments was calculated after nuclei segmentation and spot detection as described above. Only RNA foci within the detected nuclei and with intensities above a user defined threshold were considered. FISH and EdU intensities at these locations were measured as the maximum intensity in a window of +/- 300 nm.

To analyze colocalization of immunolabeled proteins and Neat1 with vDNA, we used the Costes method (Costes et al, 2004) as implemented in the coloc2 plugin of Fiji (Schindelin et al, 2012). First, nuclei were manually selected to exclude background that could lead to inappropriate thresholding during correlation analysis. For each selected region, the plugin automatically thresholds each channel such that the Pearson correlation coefficient of channel intensities below the threshold is 0. The Pearson correlation of intensities above the thresholds is then reported. Similarly to above, statistical significance was determined by comparing this correlation to that of 100 randomly jittered images, where jittering consisted in moving one channel by 3 pixels. If the Pearson correlation of the unjittered channels is positive (respectively, negative), the reported p-value reflects the proportion of jitter iterations in which the Pearson coefficient is higher (respectively, lower).

To estimate the size of nuclear vDNA foci, we used Fiji (Schindelin et al, 2012) to manually draw line profiles across EdU spots. Each intensity profile was then fitted with a Gaussian function and the corresponding full width at half maximum (FWHM) was used as a measure of the vDNA cluster size.

## Notes

### Competing Interest Statement

The authors have declared no competing interest.

### Summary of Updates

The paper now includes additionally results, notably in primary macrophages.

