## Supplementary material for "Clustering and reverse transcription of HIV-1 genomes in nuclear niches of macrophages": Supplemetary text and figures

**for**

**Rensen et al.**

**Content**

**Figure S1: Time course of HIV-1 early steps of infection in ThP1 cells**	**3**

**Figure S2: EdU-labeled HIV genomes form large nuclear foci in macrophages**	**4**

**Figure S3: In HIV-1 infected cells, EdU colocalizes with integrase (IN) and capsid (CA).**	**5**

**Figure S4: Infected macrophages display nuclear vDNA foci in absence of VPX**	**6**

**Figure S5: RNA FISH specifically labels vRNA**	**7**

**Figure S6: vDNA foci are also foci of vRNA**	**8**

**Figure S7: RNA-FISH specifically labels HIV1 RNA**	**9**

**Figure S8: Nuclear vDNA foci have lower densities of cellular DNA**	**10**

**Figure S9: Viral clusters colocalize with CPSF6**	**11**

**Figure S10: Viral clusters localize in the proximity of paraspeckle marker NEAT1**	**12**

**Figure S11: Viral clusters colocalize with the speckle marker SC35**	**13**

**Figure S12: Nuclear vDNA/vRNA clusters can form in absence of integration**	**14**

**Figure S13: Nuclear vDNA/vRNA foci form despite pharmacological inhibition of viral integration**	**15**

**Figure S14: Effect of reverse transcription inhibition on vDNA/vRNA foci**	**16**

**Figure S15: Genomic vRNA is also present in nuclei of untreated infected ThP1 cells**	**17**

**Figure S16: Effect of reverse transcription recovery after inhibition on vDNA/vRNA foci**	**18**

**Figure S17: Effect of reverse transcription recovery with nuclear import block on vDNA/vRNA foci**	**19**

**Figure S18: PF74 blocks nuclear import of vRNA**	**20**

**Figure S19: Effect of reverse transcription recovery with protracted nuclear import block on vDNA/vRNA foci**	**21**

**Figure S20: qPCR analysis of 2LTR and ALU-PCR**	**22**

**Figure S21: Nuclear RT can lead to transcription competent vDNA**	**24**

**Figure S22: Characterization of macrophages from human donor**	**25**

**Figure S23: vRNA and vDNA clusters in primary macrophages**	**27**

**Note S1: Reagents and antibodies**	**28**

**Table S1. RNA-FISH probes against HIV-1 POL gene**	**29**

**Table S2. RNA-FISH probes against GFP cDNA**	**30**

**Table S3. RNA FISH probes against LUC cDNA**	**31**

**Supplementary references:**	**32**

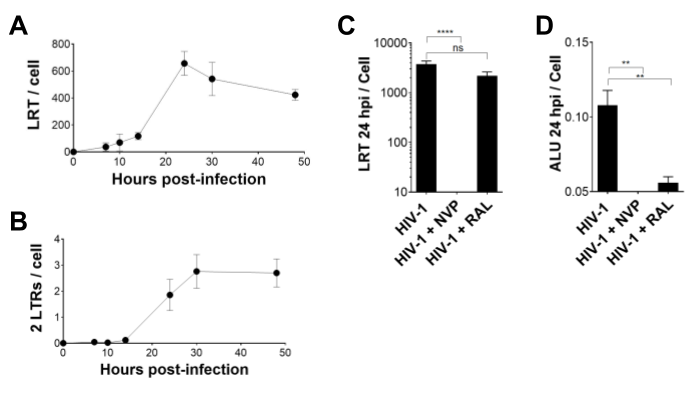

### Figure S1: Time course of HIV-1 early steps of infection in ThP1 cells

(**A**) DNA synthesis measured by qPCR of late reverse transcripts (LRT), normalized by actin.

(**B**) Formation of two long terminal repeats (2LTRs), exclusively nuclear forms of vDNA, by qPCR normalized by actin.

(**C**) DNA synthesis at 24 h post-infection measured as in **A** for HIV-1 infected cells that are untreated or treated with Nevirapine (NVP) or Raltegravir (RAL).

(**D**) Integration measured by ALU PCR, normalized by actin.

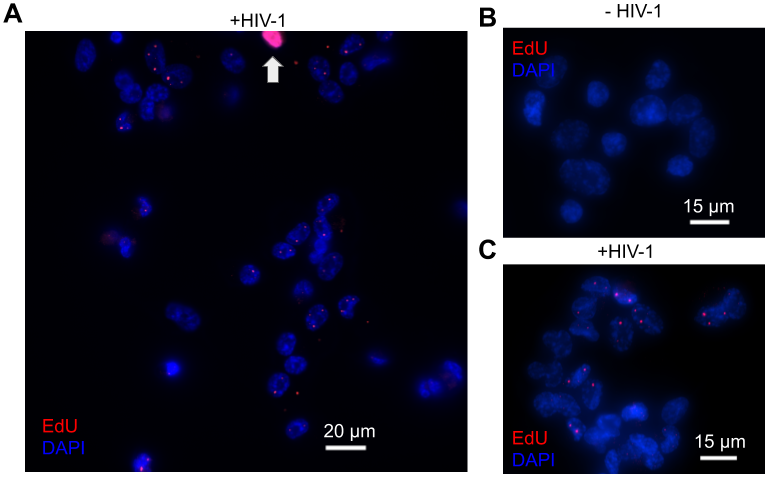

### Figure S2: EdU-labeled HIV genomes form large nuclear foci in macrophages

(**A-C**) Images of EdU labeled Thp1 cells, with EdU shown in red and DAPI in blue. (**A,C**) EdU-labeled viral DNA forms bright foci in ThP1 cells infected by HIV-1 at 48 h p.i. A small fraction of ThP1 cells displayed a very bright and uniform nuclear EdU signal, indicating that they are not differentiated (white arrow in **A**). (**B**) Non-infected cells.

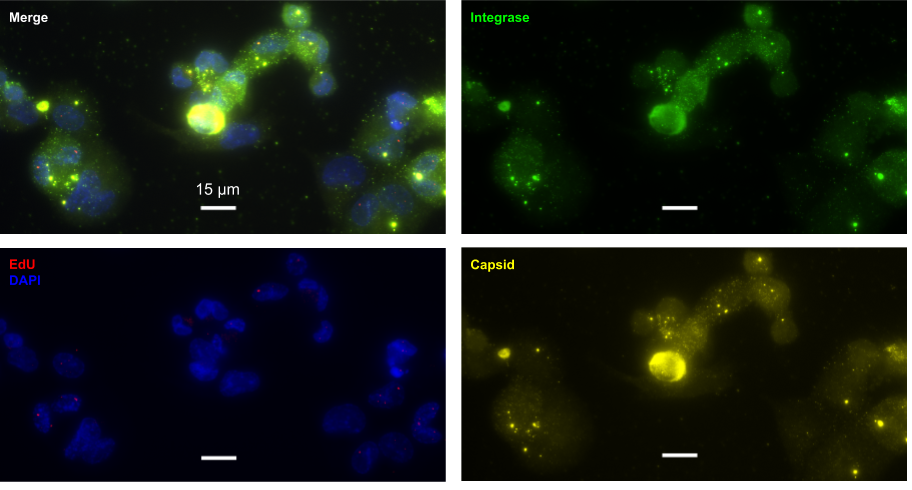

### Figure S3: In HIV-1 infected cells, EdU colocalizes with integrase (IN) and capsid (CA).

Multi-color image of HIV-1 infected ThP1 cells showing EdU (red) with immunolabeled CA (yellow) and integrase (green).

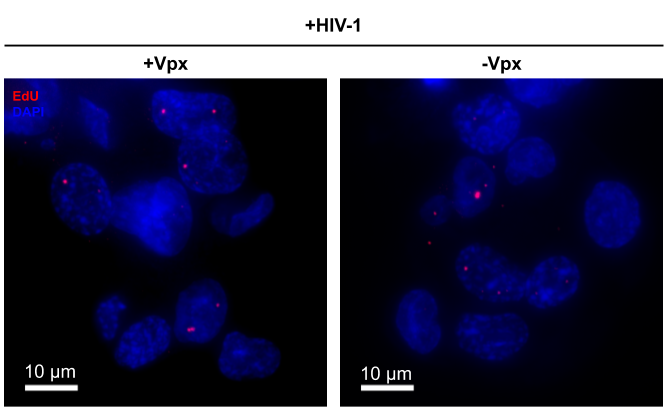

### Figure S4: Infected macrophages display nuclear vDNA foci in absence of VPX

Images of infected ThP1 cells in presence (left) or absence (right) of Vpx. Nuclear EdU foci are visible in both cases, indicating that these structures are not due to the incorporation of Vpx by HIV-1 viral particles.

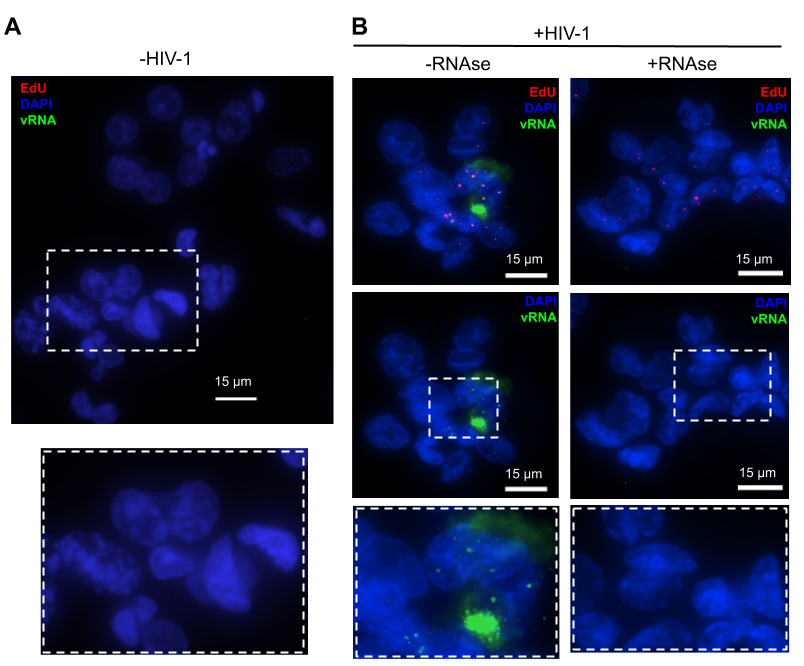

### Figure S5: RNA FISH specifically labels vRNA

Three color images of ThP1 cells with RNA-FISH against HIV-1 POL in green, EdU in red, and DAPI in blue. (**A**) In uninfected cells, no RNA-FISH signal is visible. (**B**) In cells infected with HIV-1 and not treated with RNAse (left), RNA-FISH signal is clearly visible and colocalizes with EdU in the nucleus. In infected cells treated with RNAse, EdU foci are visible but no RNA-FISH signal is visible. Only the vRNA and DAPI channels are shown in the second row for clarity. These controls demonstrate that vRNA labeling by RNA-FISH is specific. Dashed white rectangles are shown magnified in the bottom row.

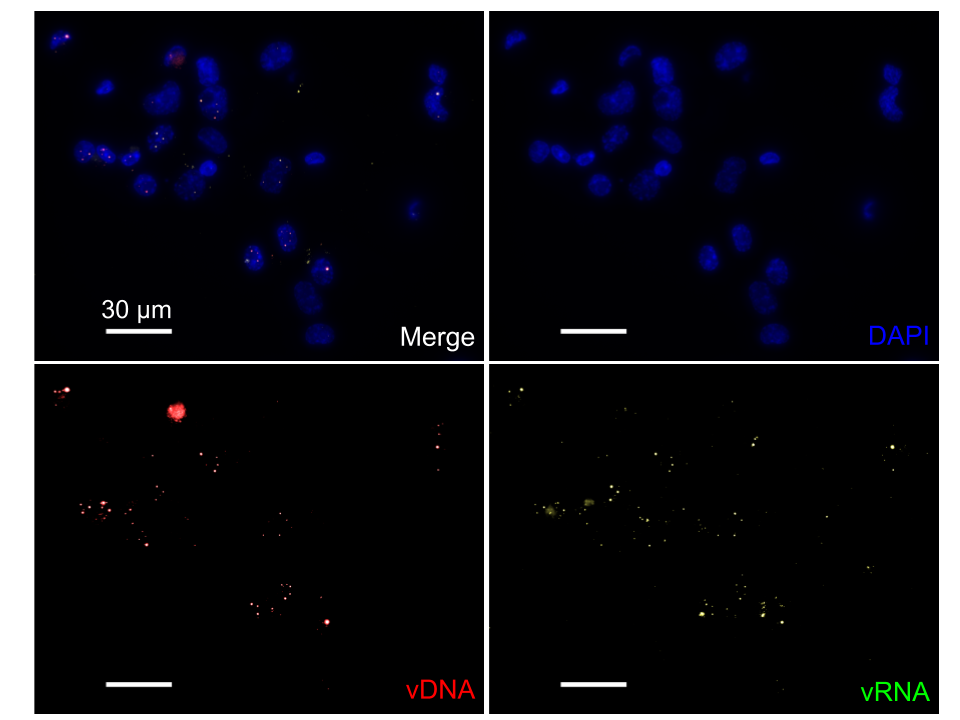

### Figure S6: vDNA foci are also foci of vRNA

Dual-color image of infected ThP1 cells at 48 h p.i. showing the viral DNA (red) and the viral RNA visualized by RNA-FISH (green). The brightness of the two bottom panels was increased for clarity.

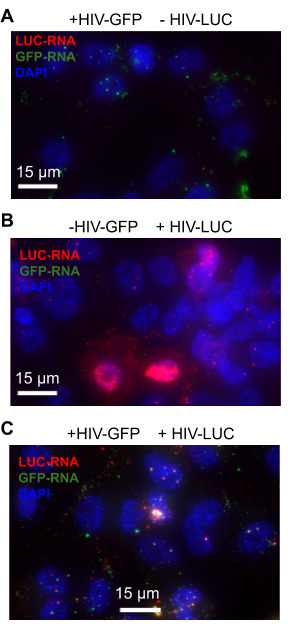

### Figure S7: RNA-FISH specifically labels HIV1 RNA

Multicolor RNA-FISH images of ThP1 cells, with FISH probes targeting LUC RNA shown in red, and FISH probes targeting GFP RNA in green. DAPI is shown in blue. (**A,B**) Control cells infected with a single viral strand to assay specificity. (**A**) Cells infected with HIV1-GFP virus only. Only green signal is visible, as expected. (**B**) Cells infected with HIV1-LUC virus only. Only red signal is visible, as expected. These images show that RNA-FISH probes are specific for GFP and LUC. **c**) Cells infected with both HIV1-GFP and HIV1-LUC virus. Both red and green signal is visible, with strong colocalization in nuclear foci.

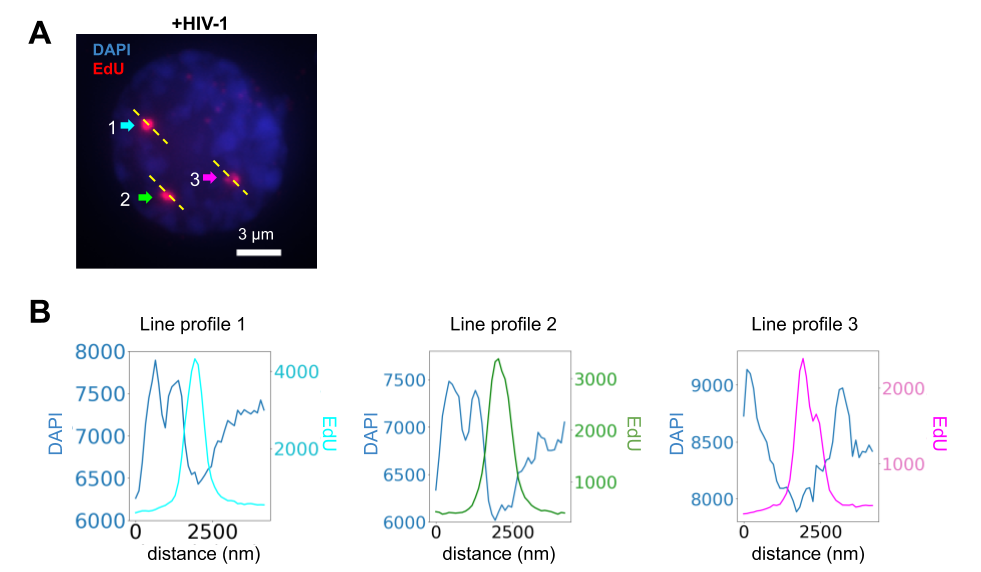

### Figure S8: Nuclear vDNA foci have lower densities of cellular DNA

(**A**) Dual color image of an infected ThP1 cell nucleus with EdU shown in red and DAPI in bue (same image as in **Fig. 1** **A**). Dashed lines are used to measure intensities of DAPI and EdU across three vDNA foci. (**B**) Intensities of DAPI and EdU along the three profiles. EdU intensity peaks coincide with local minima of DAPI intensity.

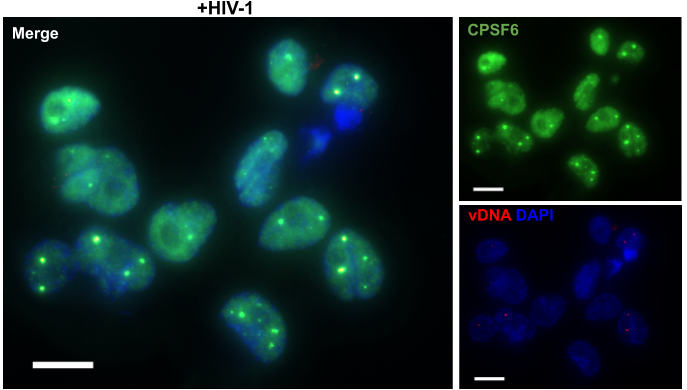

### Figure S9: Viral clusters colocalize with CPSF6

Multicolor image of infected ThP1 cells showing the vDNA (EdU) in red, the nucleus (DAPI) in blue, and immunostained CPSF6 in green. The green channel is shown separate from the blue and red channels in the smaller views on the right.

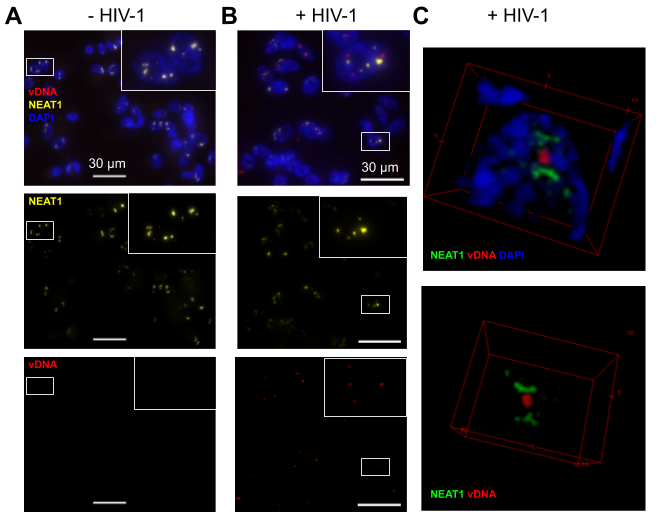

### Figure S10: Viral clusters localize in the proximity of paraspeckle marker NEAT1

**(A,B**) Multicolor images showing the immunostained long non-coding RNA NEAT1 in yellow, vDNA (EdU) in red, and the nucleus (DAPI) in blue in uninfected (**A**) or infected (**B**) ThP1 cells. Note that NEAT1 accumulates in nuclear foci, as expected for speckles, and that vDNA foci are visible in close proximity to NEAT foci, but do not colocalize with NEAT1. (**C**) 3D renderings of confocal images of infected ThP1 cells showing NEAT1 in green, vDNA (EdU) in red, and the nucleus (DAPI) in blue.

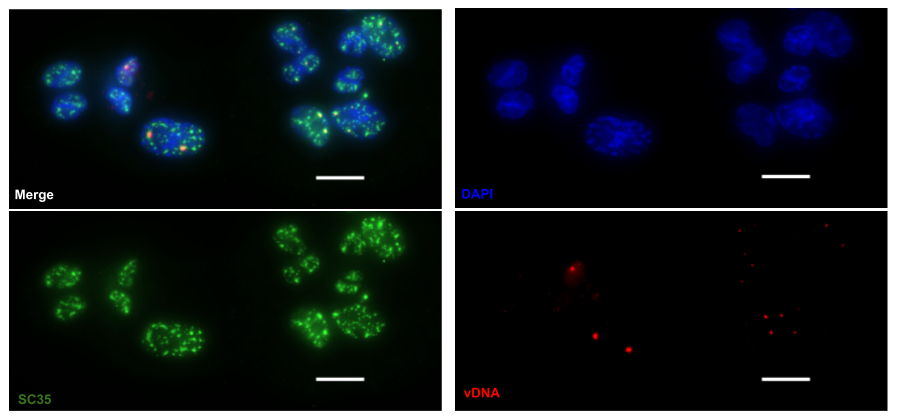

### Figure S11: Viral clusters colocalize with the speckle marker SC35

Multicolor image of infected ThP1 cells showing immunostained SC35 in green, the vDNA (EdU) in red, the nucleus (DAPI) in blue. vDNA foci are observed in a subset of SC35 foci.

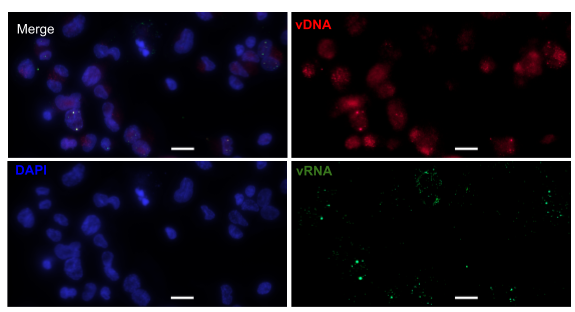

### Figure S12: Nuclear vDNA/vRNA clusters can form in absence of integration

Multicolor image of ThP1 cells infected with an integration-deficient HIV-1 carrying the mutation D116A in the catalytic site of integrase (IN). The EdU-labeled vDNA is shown in red and the vRNA detected by RNA-FISH in green. The nucleus (DAPI) is shown in blue. The brightness of the vRNA image (bottom right) was increased for better visibility.

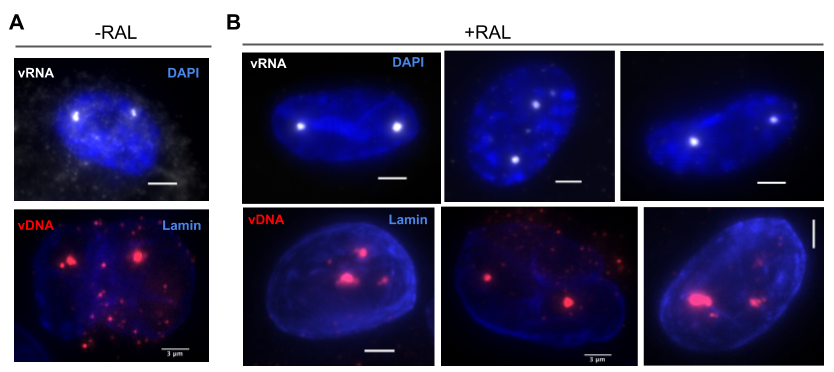

### Figure S13: Nuclear vDNA/vRNA foci form despite pharmacological inhibition of viral integration

Images of infected ThP1 cells treated with 10 uM Raltegravir (RAL), an inhibitor of integration (**B**) and infected control cells without RAL (**A**). The EdU-labeled vDNA is shown in red and the vRNA detected by RNA-FISH in white. The nucleus (DAPI or lamin) is shown in blue. Clear vDNA/vRNA foci are visible both in untreated cells and RAL-treated cells.

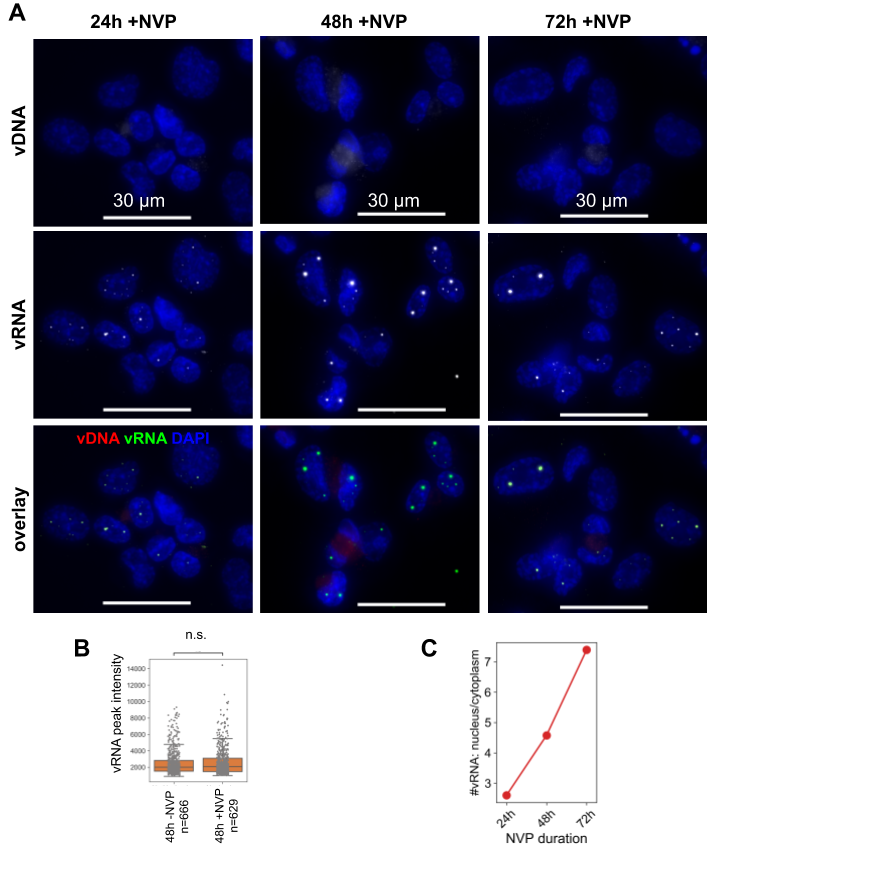

### Figure S14: Effect of reverse transcription inhibition on vDNA/vRNA foci

(**A**) Multicolor images of infected ThP1 cells treated with Nevirapine (NVP) for 24 h, 48 h or 72 h p.i., with labeled vDNA (EdU), vRNA (RNA-FISH), and nuclei (DAPI). (**B**) Peak intensities of vRNA at 48 h p.i. in cells treated with NVP (+NVP) or untreated (-NVP). The intensity distributions do not differ significantly (Wilcoxon test: p=0.97). (**C**) Ratio of nuclear to cytoplasmic vRNA as function of NVP treatment duration. Quantifications were made with FISH-Quant and Imjoy (Mueller *et al*, 2013; Ouyang *et al*, 2019).

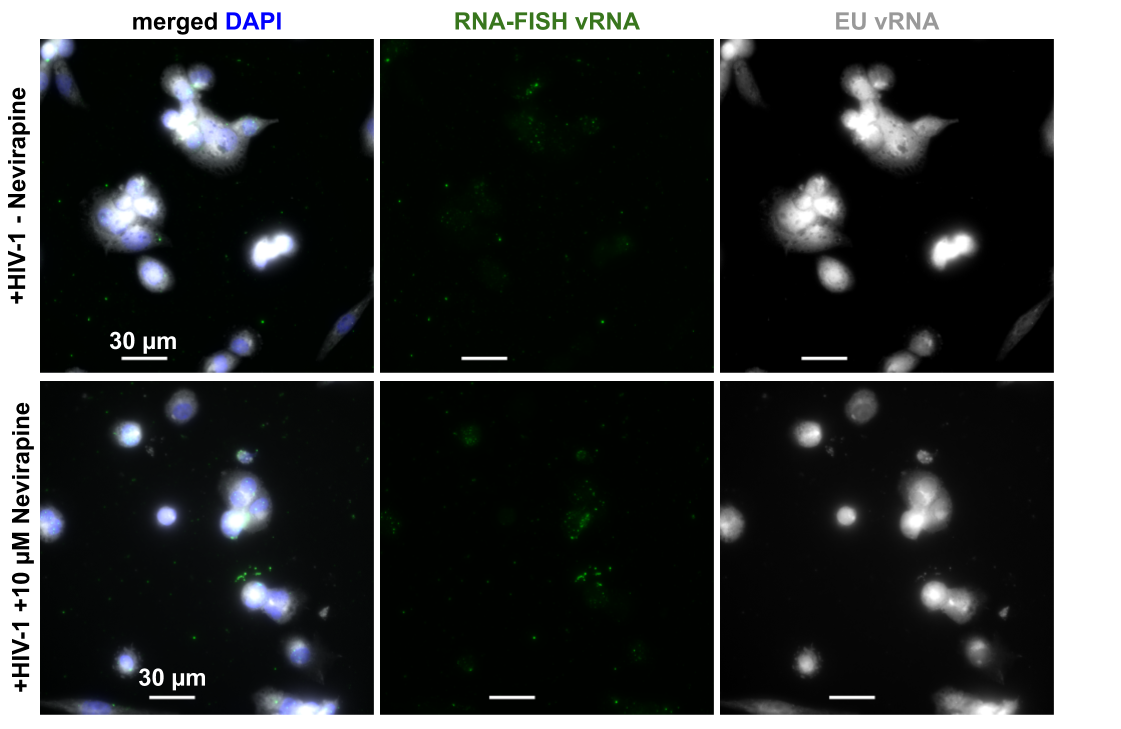

### Figure S15: Genomic vRNA is also present in nuclei of untreated infected ThP1 cells

Multicolor images of ThP1 cells infected with an EU-labeled virus, in untreated cells (top) and cells exposed to Nevirapine (bottom). The RNA-FISH and EU images are shown separately in green and white, respectively. The left column shows merged images with DAPI and blue, RNA-FISH in green, and EU in white.

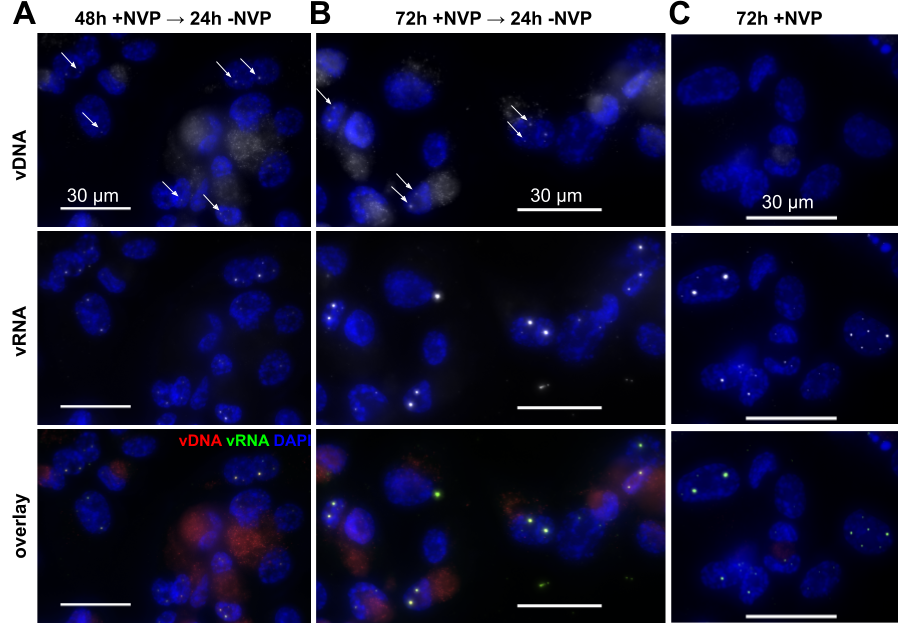

### Figure S16: Effect of reverse transcription recovery after inhibition on vDNA/vRNA foci

Multicolor images of infected ThP1 cells with labeled vDNA (EdU), vRNA (RNA-FISH), and nuclei (DAPI). (**A,B**) Cells were first treated with Nevirapine (NVP) for 48 h (left) or 72 h p.i. (right), then the drug was washed out and cells were fixed for imaging 24 h later. Arrows point to examples of vDNA foci. (**C**) Cells treated with NVP for 48 h p.i. and fixed thereafter, for comparison; these images are identical to **Fig. S17 A** (middle column). No vDNA foci are seen.

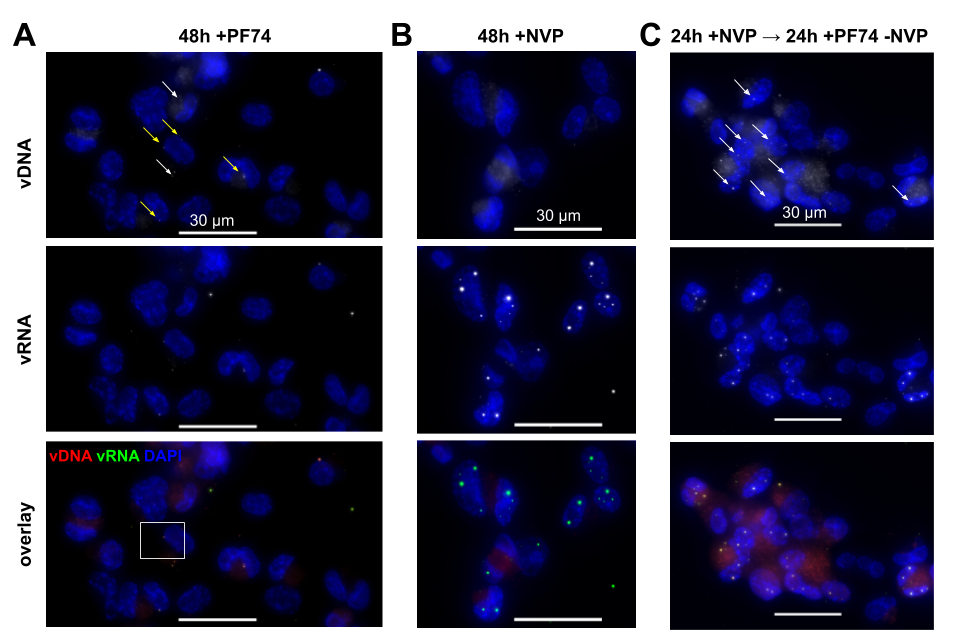

### Figure S17: Effect of reverse transcription recovery with nuclear import block on vDNA/vRNA foci

Multicolor images of infected ThP1 cells with labeled vDNA (EdU), vRNA (RNA-FISH), and nuclei (DAPI). (**A**) Cells were exposed to the drug PF74 for 48 h p.i. Isolated vDNA  and vRNA spots can be seen in the cytoplasm and at the nuclear envelope (white and yellow arrows, respectively), but not inside nuclei, confirming efficient block of nuclear HIV-1 import by PF74. (**B**) for comparison, cells exposed to the reverse transcription inhibitor NVP for 48 h p.i. display no detectable vDNA foci, but show large vRNA foci; images are identical to **Fig. S17 A**. (**C**) Cells were exposed to NVP for 24 h p.i., then NVP was washed out and replaced by PF74, and cells were fixed for imaging 24 h later. vDNA foci colocalizing with vRNA foci can be seen inside nuclei (arrows).

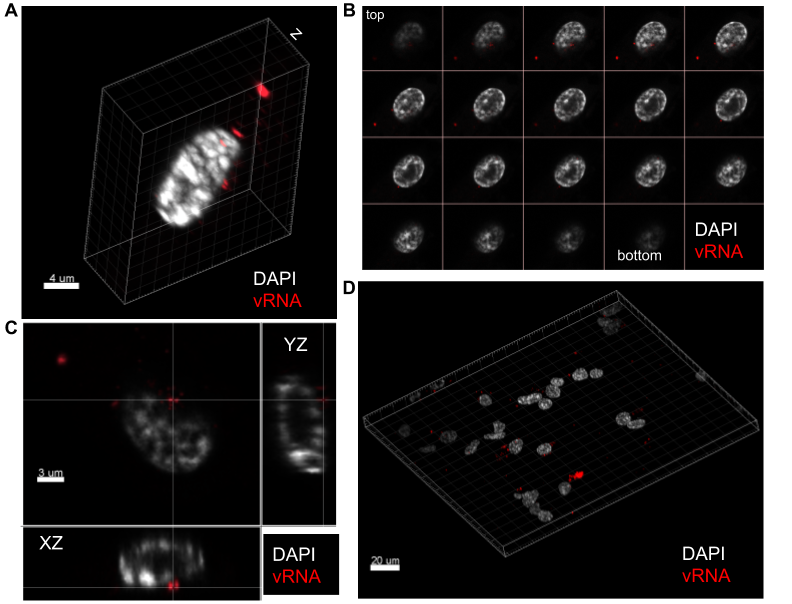

### Figure S18: PF74 blocks nuclear import of vRNA

3D confocal z-stack images of PF74 treated cells at 48 h p.i. To assess if PF74 efficiently blocks nuclear import in infected ThP1 cells, vRNA was labelled by RNA-FISH (red) and nuclei stained with DAPI (white). (**A,B**) Perspective view (**A**) and montage view (**B**) of a single cell. (**C**) Orthogonal views of the z-stack, where the central image shows an XY slice, and images to the right and below show XZ and YZ slices at positions indicated by the lines. (**D**) Larger image, showing more than 20 cells. In all images, vRNA can be seen in the cellular cytoplasm or close to the nucleus, but is absent from the nucleus, confirming that PF74 efficiently blocks HIV-1 nuclear import in ThP1 cells.
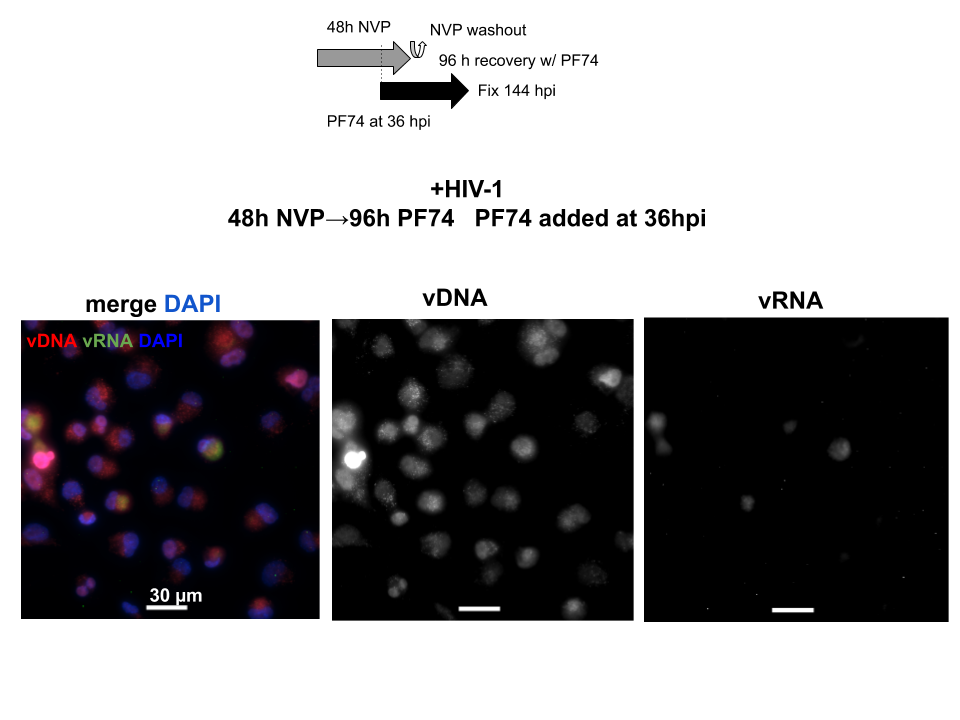

### Figure S19: Effect of reverse transcription recovery with protracted nuclear import block on vDNA/vRNA foci

Multicolor images of infected ThP1 cells with labeled vDNA (EdU), vRNA (RNA-FISH), and nuclei (DAPI). Cells were exposed to the reverse transcription inhibitor NVP for 48 h p.i. At 36 h p.i., 1.5 µM of PF74, which blocks nuclear import, was added to the cells along with NVP. After NVP washout, the cells were continuously exposed to PF74 for an additional 96 h until fixation. vDNA foci colocalizing with vRNA foci can be seen inside nuclei.

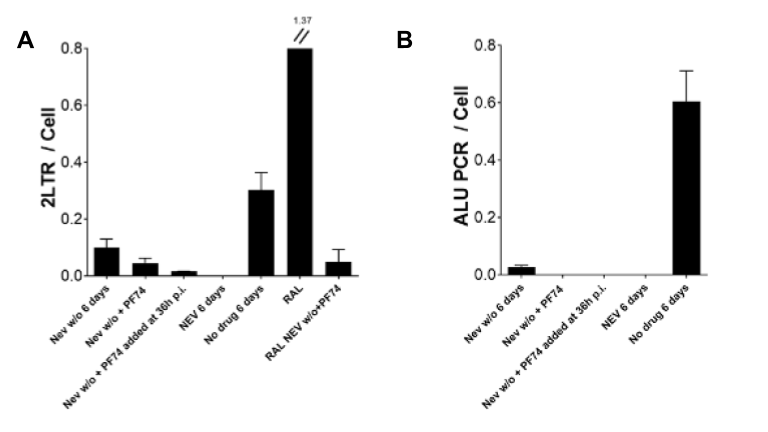

### Figure S20: qPCR analysis of 2LTR and ALU-PCR

Samples treated and untreated with drugs have been analyzed by qPCR to amplify episomal nuclear vDNA forms, 2LTRs (**A**) and normalized by actin. The number of provirus per cell has been analyzed by ALU PCR and normalized by actin, ALU values were normalized considering respective controls consisting in samples treated with RAL (**B**).

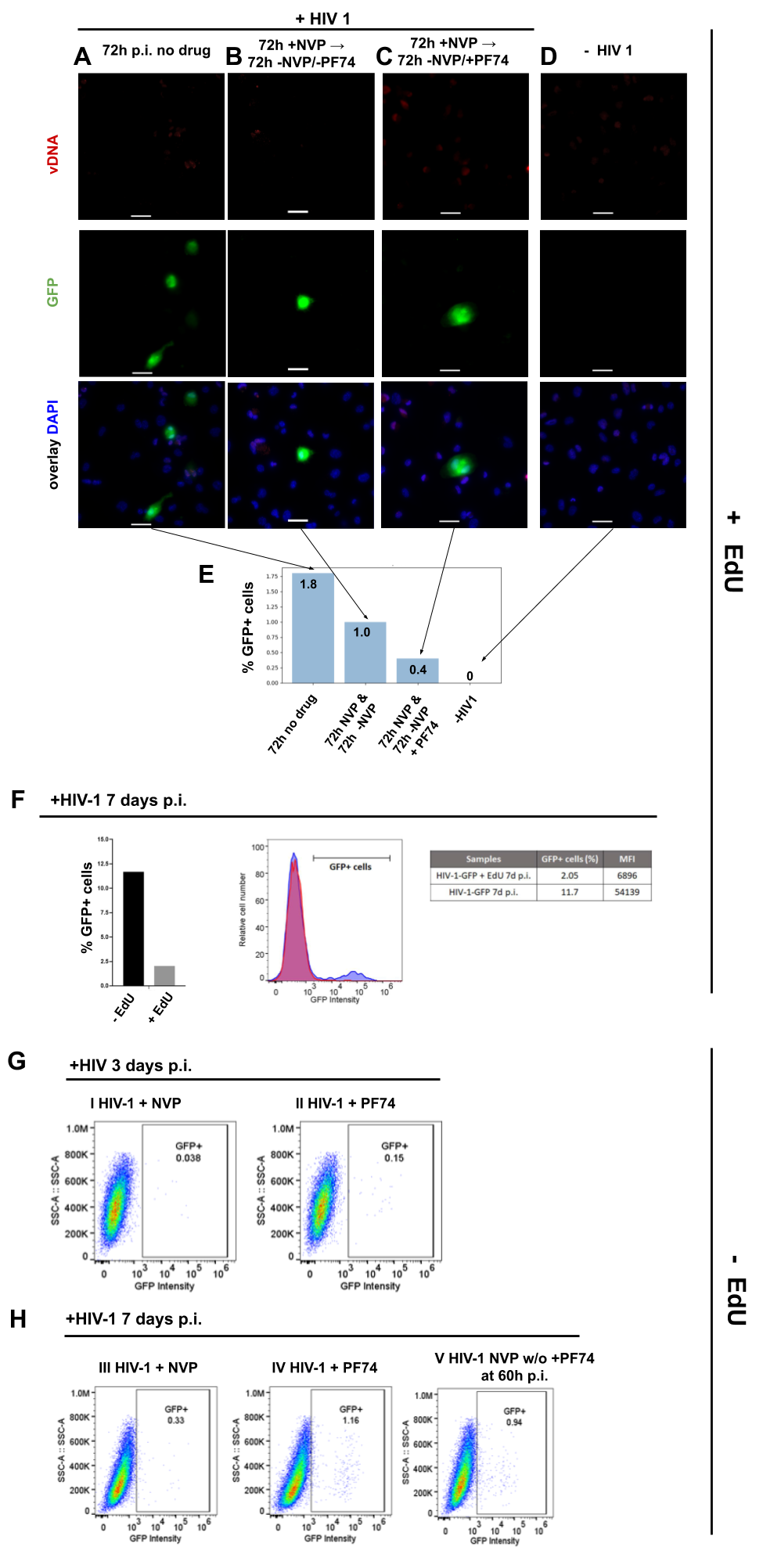

**Figure S21:** Legend on next page

### Figure S21: Nuclear RT can lead to transcription competent vDNA

(**A-D**) Multicolor images of ThP1 cells infected with a GFP-reporter virus. vDNA (EdU) is labeled in red, GFP in green and nuclei (DAPI) in blue. (**A**) Untreated, infected ThP1 cells fixed at 3 d (72 h) p.i. We chose this time point for comparison with the three other conditions **B-D** in order to allow for the same duration of RT before fixation (3d). (**B**) Infected cells were exposed to NVP for 72 h, then NVP was washed out and cells were cultured for another 72 h and fixed at 6 d p.i. (**C**) Infected cells were cultured for 3d p.i., then NVP was washed out and cells were exposed to the nuclear import inhibitor PF74 for another 3d and fixed at 6 d p.i.  (**D**) Non infected control cells. (**E**) Percentage of GFP positive cells detected in the four experimental conditions **A-D**. (**F**) EdU incorporation affects transcription. Left: FACS analysis of GFP positive cells in infected cells at 7 d p.i. with (grey) or without (black) incubation with EdU. (**G,H**) Percentage of GFP positive cells at 3 d (**G**) and 7 d (**H**) post infection analyzed by FACS, in absence of EdU. (**G**) Panel **I**: Infected cells were exposed to NVP and fixed at 3 d p.i. Panel **II**: Infected cells were exposed to 1.5 µM PF74 and fixed at 3 d p.i. (**H**) Panel **III**: Infected cells were exposed to NVP and fixed at 7 d p.i. Panel **IV**: Infected cells were exposed to 1.5 µM PF74 and fixed at 7d pi. Panel **V**: Cells were exposed to NVP for 3 d p.i. At 60 h p.i., 1.5 µM of PF74 was added along with NVP. After NVP washout, the cells were continuously exposed to PF74 for an additional 4 d until fixation.

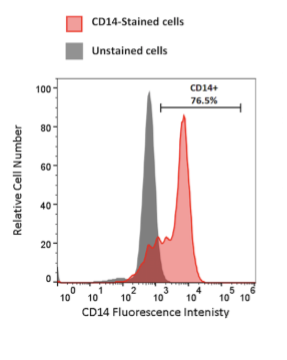

### Figure S22: Characterization of macrophages from human donor

Monocyte derived macrophages (MDMs) from donor 2 were differentiated and cultured in RPMI complemented with MCSF and 10% foetal calf serum. FACS analysis of CD14 was done at 7 days p.i., at the same moment of infectivity analysis.

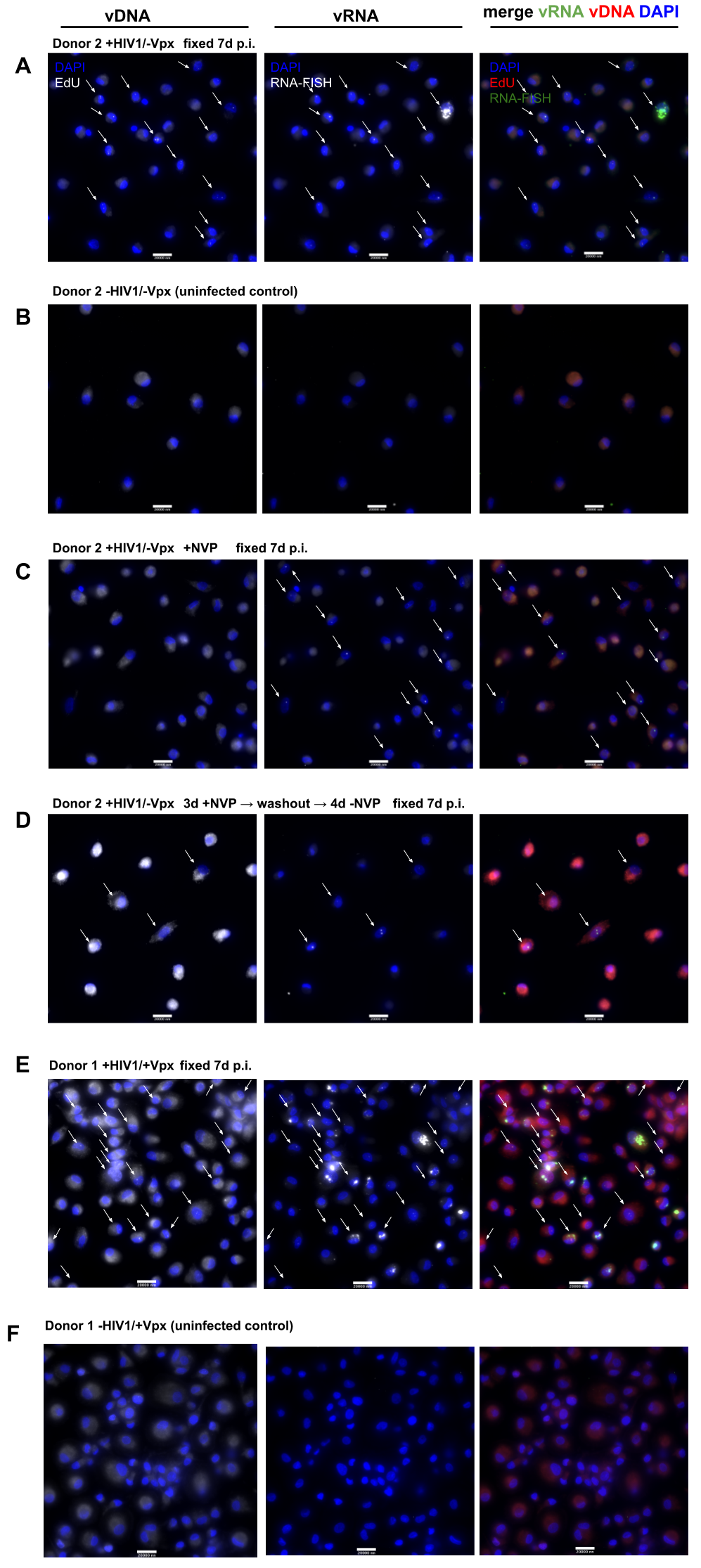

**Figure S23:** Legend on next page

### Figure S23: vRNA and vDNA clusters in primary macrophages

Multicolor images of MDMs from donors 1 and 2, showing DAPI together with vDNA (EdU), vRNA (RNA-FISH), or both. Cells in (**A,C,D**) were infected with HIV-1 (without Vpx) and fixed for imaging 6 d p.i. Cells in (**B**) were uninfected. **C**) cells were treated with Nevirapine (NVP) during 6 d. **D**) Cells were treated with NVP for 3 d, then NVP was washed out and fixed 4 d later, i.e. at 7 d p.i. Cells from donor 1 (**E,F**) were infected with HIV-1 (with Vpx) (**E**) or left uninfected (**F**) and fixed for imaging 7 d p.i..

### Note S1: Reagents and antibodies

Primary antibodies

Anti-p24 antibody NIH183-H12-5C (NIH reagent, IF dil. 1:400)

Anti-HA high-affinity antibody (11867423001) Roche (IF dil. 1:500)

Anti NONO (C-terminal) antibody, Sigma N8664 (IF dil. 1:400)

Anti-CPSF6 antibody, NBP1-85676, Novus (IF dil. 1:500)

Anti-SC35 antibody, Abcam ab11826 (IF dil. 1:200)

Anti-A/C lamin (636) antibody, Santa Cruz Biotechnology sc-7292 (IF dil: 1:200)

Secondary antibodies

Goat anti-mouse Alexa Fluor Plus 488 (A32723)

Goat anti-rat Alexa 647 (A21247) Thermofisher scientific.

Kits and reagents

FISH probes, human NEAT1 5' segment with QUASAR 570 DYE (Stellaris SMF-2036-1)

Click-chemistry kit EdU (Click-iT™ Plus Alexa Fluor™ 647 Picolyl Azide Toolkit, Invitrogen™, C10643)

Click-iT® RNA Imaging Kit (Click-iT™ RNA Alexa Fluor™ 488 Imaging Kit, Invitrogen™, C10329)

Nevirapine (Sigma SML0097)

Raltegravir (Merck CDS023737-25MG)

PF74 (Sigma SML0835)

Phorbol-12-myristate 13-acetate (Sigma-Aldrich, Saint Luis, MO, USA)

Paraformaldehyde (32% Paraformaldehyde (formaldehyde) aqueous solution, Electron Microscopy Sciences)

ProLong Gold antifade mounting medium (Molecular Probes)

### Table S1. RNA-FISH probes against HIV-1 POL gene

| **Probe Name** | **Sequences** |
| --- | --- |
| POL-01 | GGG GAT TGT AGG GAA TTC CAA ATT CCT GCT TTT ACA CTC GGA CCT CGT CGA CAT GCA TT |
| POL -02 | CTT TTA GCT GAC ATT TAT CAC AGC TGG CTA TTA CAC TCG GAC CTC GTC GAC ATG CAT T |
| POL -03 | GTG TGC TGG TAC CCA TGC CAG ATA GAC TTA CAC TCG GAC CTC GTC GAC ATG CAT T |
| POL -04 | AAT ACT GGA GTA TTG TAT GGA TTT TCA GGC CCT TAC ACT CGG ACC TCG TCG ACA TGC ATT |
| POL -05 | TTT TAC TGG TAC AGT CTC AAT AGG GCT AAT GGT TAC ACT CGG ACC TCG TCG ACA TGC ATT |
| POL -06 | TAT GTT GAC AGG TGT AGG TCC TAC TAA TAC TGT TAC ACT CGG ACC TCG TCG ACA TGC ATT |
| POL -07 | CTA ATC CTC ATC CTG TCT ACT TGC CAT TAC ACT CGG ACC TCG TCG ACA TGC ATT |
| POL -08 | CAA TCA TCA CCT GCC ATC TGT TTT CCA TTT ACA CTC GGA CCT CGT CGA CAT GCA TT |
| POL -09 | TTT CCA AAG TGG ATT TCT GCT GTC CCT GTA TTA CAC TCG GAC CTC GTC GAC ATG CAT T |
| POL -10 | TTG TGG ATG AAT ACT GCC ATT TGT ACT GCT GTT ACA CTC GGA CCT CGT CGA CAT GCA TT |
| POL -11 | TTA AGA TGT TCA GCC TGA TCT CTT ACC TGT TTA CAC TCG GAC CTC GTC GAC ATG CAT T |
| POL -12 | TAC AGT CTA CTT GTC CAT GCA TGG CTT CTT ACA CTC GGA CCT CGT CGA CAT GCA TT |
| POL -13 | TCA TGT TCA TCT TGG GCC TTA TCT ATT CCT TAC ACT CGG ACC TCG TCG ACA TGC ATT |
| POL -14 | TGT CAG TTA GGG TGA CAA CTT TTT GTC TTC CTT TAC ACT CGG ACC TCG TCG ACA TGC ATT |
| POL -15 | TGC TCC TAC TAT GGG TTC TTT CTC TAA CTT TAC ACT CGG ACC TCG TCG ACA TGC ATT |
| POL -16 | TCT GTT AGT GCT TTG GTT CCT CTA AGG AGT TTT TAC ACT CGG ACC TCG TCG ACA TGC ATT |
| POL -17 | CTG TAT GTC ATT GAC AGT CCA GCT GTC TTT TTT ACA CTC GGA CCT CGT CGA CAT GCA TT |
| POL -18 | TGG CAG CAC TAT AGG CTG TAC TGT CCT TAC ACT CGG ACC TCG TCG ACA TGC ATT |
| POL -19 | TCT GAT GTT TTT TGT CTG GTG TGG TAA GTC CCT TAC ACT CGG ACC TCG TCG ACA TGC ATT |
| POL -20 | CCT CAA CAG ATG TTG TCT CAG CTC CTC TTA CAC TCG GAC CTC GTC GAC ATG CAT T |
| POL -21 | ATT GCT GGT GAT CCT TTC CAT CCC TGT TAC ACT CGG ACC TCG TCG ACA TGC ATT |
| POL -22 | TTT CTT TTT TAA CCC TGC GGG ATG TGG TAT TCT TAC ACT CGG ACC TCG TCG ACA TGC ATT |
| POL -23 | TTT AAC TTT TGG GCC ATC CAT TCC TGG CTT ACA CTC GGA CCT CGT CGA CAT GCA TT |
| POL -24 | CCC TAT CTT TAT TGT GAC GAG GGG TCG TTG TTA CAC TCG GAC CTC GTC GAC ATG CAT T |

### Table S2. RNA-FISH probes against GFP cDNA

| **Probe Name** | **Sequences** |
| --- | --- |
| GFP-01 | TTC AGC TCG ATG CGG TTC ACC AGG GTT TAC ACT CGG ACC TCG TCG ACA TGC ATT |
| GFP-02 | GAA GAT GGT GCG CTC CTG GAC GTA GCT TAC ACT CGG ACC TCG TCG ACA TGC ATT |
| GFP-03 | AGT CGT GCT GCT TCA TGT GGT CGG GGT TAC ACT CGG ACC TCG TCG ACA TGC ATT |
| GFP-04 | GCG GCT GAA GCA CTG CAC GCC GTT TAC ACT CGG ACC TCG TCG ACA TGC ATT |
| GFP-05 | GTC AGG GTG GTC ACG AGG GTG GGC CAT TAC ACT CGG ACC TCG TCG ACA TGC ATT |
| GFP -06 | TTC AGG GTC AGC TTG CCG TAG GTG GCT TAC ACT CGG ACC TCG TCG ACA TGC ATT |
| GFP -07 | TTG TGG CCG TTT ACG TCG CCG TCC AGT TAC ACT CGG ACC TCG TCG ACA TGC ATT |
| GFP -08 | GAA CAG CTC CTC GCC CTT GCT CAT TAC ACT CGG ACC TCG TCG ACA TGC ATT |
| GFP -09 | CGC TGC CGT CCT CGA TGT TGT GGC TTA CAC TCG GAC CTC GTC GAC ATG CAT T |
| GFP -10 | GTT GCC GTC CTC CTT GAA GTC GAT GCT TAC ACT CGG ACC TCG TCG ACA TGC ATT |
| GFP -11 | CGC CCT CGA ACT TCA CCT CGG CGT TAC ACT CGG ACC TCG TCG ACA TGC ATT |
| GFP -12 | TGA ACT TGT GGC CGT TTA CGT TAC ACT CGG ACC TCG TCG ACA TGC ATT |
| GFP -13 | TGG TGC AGA TGA ACT TCA GGT TAC ACT CGG ACC TCG TCG ACA TGC ATT |
| GFP -14 | TAG GTC AGG GTG GTC ACG AGT TAC ACT CGG ACC TCG TCG ACA TGC ATT |
| GFP -15 | TGG CGG ACT TGA AGA AGT CGT TAC ACT CGG ACC TCG TCG ACA TGC ATT |
| GFP -16 | CTT GAA GAA GAT GGT GCG CTT TAC ACT CGG ACC TCG TCG ACA TGC ATT |
| GFP -17 | TTG AAG TCG ATG CCC TTC AGT TAC ACT CGG ACC TCG TCG ACA TGC ATT |
| GFP -18 | CTG TTG TAG TTG TAC TCC AGT TAC ACT CGG ACC TCG TCG ACA TGC ATT |
| GFP -19 | TTG TCG GCC ATG ATA TAG ACT TAC ACT CGG ACC TCG TCG ACA TGC ATT |
| GFP -20 | CTT GAA GTT CAC CTT GAT GCT TAC ACT CGG ACC TCG TCG ACA TGC ATT |
| GFP -21 | TGC TCA GGT AGT GGT TGT CGT TAC ACT CGG ACC TCG TCG ACA TGC ATT |
| GFP -22 | GTC ACG AAC TCC AGC AGG ACT TAC ACT CGG ACC TCG TCG ACA TGC ATT |
| GFP -23 | TAC TTG TAC AGC TCG TCC ATT TAC ACT CGG ACC TCG TCG ACA TGC ATT |

### Table S3. RNA FISH probes against LUC cDNA

| **Probe Name** | **Sequences** |
| --- | --- |
| LUC-01 | CCT GAT AGC CTT TGT ACT TAA TCA GAG ACT TCT TAC ACT CGG ACC TCG TCG ACA TGC ATT |
| LUC -02 | CAC ACA CAG TTC GCC TCT TTG ATT AAC GCT TAC ACT CGG ACC TCG TCG ACA TGC ATT |
| LUC -03 | AGC GTT TTC CCG GTA TCC AGA TCC ACA TTA CAC TCG GAC CTC GTC GAC ATG CAT T |
| LUC -04 | CAG AAT AGC TGA TGT AGT CTC AGT GAG CCC TTA CAC TCG GAC CTC GTC GAC ATG CAT T |
| LUC -05 | ATC AGT GCA ATT GTC TTG TCC CTA TCG AAG TTA CAC TCG GAC CTC GTC GAC ATG CAT T |
| LUC -06 | ACA TCG ACT GAA ATC CCT GGT AAT CCG TTT TTA CAC TCG GAC CTC GTC GAC ATG CAT T |
| LUC -07 | TTA CAC GGC GAT CTT TCC GCC CTT CTT TAC ACT CGG ACC TCG TCG ACA TGC ATT |
| LUC -08 | TCG AGT TTT CCG GTA AGA CCT TTC GGT ACT TCT TAC ACT CGG ACC TCG TCG ACA TGC ATT |
| LUC -09 | TTT CGC GGT TGT TAC TTG ACT GGC GAC GTA ATT ACA CTC GGA CCT CGT CGA CAT GCA TT |
| LUC -10 | CAC GAT CTC TTT TTC CGT CAT CGT CTT TCC TTA CAC TCG GAC CTC GTC GAC ATG CAT T |
| LUC -11 | GAC ACC TGC GTC GAA GAT GTT GGG GTG TTT TAC ACT CGG ACC TCG TCG ACA TGC ATT |
| LUC -12 | GCG GTC AAC GAT GAA GAA GTG TTC GTC TTC GTT TAC ACT CGG ACC TCG TCG ACA TGC ATT |
| LUC -13 | ATC CTT GTC AAT CAA GGC GTT GGT CGC TTT ACA CTC GGA CCT CGT CGA CAT GCA TT |
| LUC -14 | ATC CTT GCC TGA TAC CTG GCA GAT GGT TAC ACT CGG ACC TCG TCG ACA TGC ATT |
| LUC -15 | GGG AGC GCC ACC AGA AGC AAT TTC GTG TTA CAC TCG GAC CTC GTC GAC ATG CAT T |
| LUC -16 | ATC GTA TTT GTC AAT CAG AGT GCT TTT GGC GAT TAC ACT CGG ACC TCG TCG ACA TGC ATT |
| LUC -17 | CTT TGA ATC TTG TAA TCC TGA AGG CTC CTC TTA CAC TCG GAC CTC GTC GAC ATG CAT T |
| LUC -18 | GGT AGG CTG CGA AAT GCC CAT ACT GTT GTT ACA CTC GGA CCT CGT CGA CAT GCA TT |
| LUC -19 | ATT CAC GTT CAT TAT AAA TGT CGT TCG CGG GCT TAC ACT CGG ACC TCG TCG ACA TGC ATT |
| LUC -20 | AGT GAT GTC CAC CTC GAT ATG TGC ATC TTT ACA CTC GGA CCT CGT CGA CAT GCA TT |
| LUC -21 | CTT CAT AGC CTT ATG CAG TTG CTC TCC TTA CAC TCG GAC CTC GTC GAC ATG CAT T |
| LUC -22 | CGG TTC CAT CTT CCA GCG GAT AGA ATG GTT ACA CTC GGA CCT CGT CGA CAT GCA TT |
| LUC -23 | CCG GGC CTT TCT TTA TGT TTT TGG CGT CTT TTA CAC TCG GAC CTC GTC GAC ATG CAT T |
